# High-Dimensional Single-Cell Analysis Reveals Coordinated Age-Dependent Neuroinflammatory Microglia-T cell Circuits in the Brain

**DOI:** 10.64898/2025.12.10.693494

**Authors:** Md Akkas Ali, Md Hasanul Banna Siam, Donald Vardaman, Chase Bolding, J. Nicholas Brazell, Ashleigh D. Whatley, Christopher A. Risley, Harrison Tidwell, Syed Nakib Hossain, Juhi Samal, Ashley S. Harms, Mallikarjun Patil, Daniel J. Tyrrell

**Author notes:** Corresponding author:* Daniel J. Tyrrell, PhD, University of Alabama at Birmingham PBMR2 #504, 901 19^th^ St. S., Birmingham, AL 35205.

## Abstract

Aging and cerebrovascular pathology drive neuroinflammation in vascular dementia (VaD) but immune mechanisms underlying this interplay remain unresolved. Leveraging multi-modal high-dimensional imaging, flow cytometry, and split pool ligation transcriptomic sequencing in a mouse model of VaD, we constructed a brain immune cell atlas spanning young and aged mice in health and disease. We profiled microglia, T cells, macrophages, neutrophils, and B cells and integrated transcriptomics, cell-cell communication, multiplex imaging, and comparative analysis with human microglia. We found striking depletion of *Ccr7*^+^ naïve T cells and expansion of *Gzmk*^+^ cytotoxic *Cd8*^+^ effector memory T cells in the aging brain. At the same time, microglia shifted toward a pro-inflammatory state with enhanced activity of major histocompatibility class complex I (MHC-I) to T cell receptor and co-stimulation from CD86 to CD28. These shifts suggest enhanced neuroinflammatory polarization within the aged brain and in VaD. These signals were strongest from activated microglia to *Gzmk*^+^ *Cd8*^+^ T_EM_ cells, indicating that age-related microglial polarization may sustain cytotoxic T cell activation in the aged brain. Our findings suggest pro-inflammatory microglia and *Gzmk*^+^ CD8^+^ T_EM_ cells are central drivers of immune brain aging and highlights a therapeutic potential to disrupt age-related neuroinflammatory cascades in VaD.

## INTRODUCTION

The most common forms of dementia are Alzheimer’s disease (AD) and vascular dementia (VaD); however, nearly 80% of AD patients have vascular pathological at autopsy.^1–3^ Pure vascular dementia, without proteinopathy, affects an estimated 10 million people globally.^1–3^ Management of VaD, AD with vascular pathology, and other vascular contributions to cognitive impairment and dementia (VCID) involve management of cardiovascular risk factors to improve cerebral blood flow.^4^ Despite the prevalence of VCID and VaD in older people, our understanding of how the various cells within the brain contribute is lacking.^5^

Rodent models have been employed to better understand VaD including one which recapitulated the focal white matter ischemic lesions found in human VaD.^6–8^ Despite some advances in our knowledge of VaD progression relatively little is understood about the contribution of immune cells to disease pathogenesis. Nearly all rodent studies of VaD employ juvenile animals between 8-12 weeks of age, which is equivalent to 20-30 years of age in humans^9^; however, greater than 90% of dementia occurs after the age of 65 years.^10^ Aging is the strongest clinical predictor of dementia^11^, and aging is accompanied by significant alterations in immune cells in the brain including microglia and peripheral immune cells.^12–14^ Thus, most rodent models of VaD do not recapitulate the immune landscape present during VaD pathogenesis in humans.

A key unanswered question in VaD is how immune cells promote disease. Microglia are critical for maintaining homeostasis in the brain, yet in the context of aging, they can also promote neurodegenerative disease.^15–17^ In the aged brain, microglia exhibit a marked shift from quiescent, homeostatic functions towards inflammatory signaling that can exacerbate age-related neuropathology.^16,17^ Despite a core expression profile of disease-associated microglia, the overall diversity of microglia is highly contextualized and fine-tunes their specific functions.^15^ Broad microglial heterogeneity exists in the context of neurodegenerative disease. A key function of microglia is production of immune modifying factors such as cytokines and chemokines that can act as co-stimulatory agents. These can modify other nearby immune cells in the brain such as T cells. The mechanisms by which aging microglia interact with other immune cells to influence neurodegeneration remain poorly understood.

In blood and secondary lymphoid organs, aging is associated with a loss of naïve T cells (CD44^−^CD62L^+^) due to age-associated thymic involution and an expansion of effector memory and central memory (CD44^+^) subsets.^14,18–23^ In the brain, cytotoxic CD8^+^ T cells are either recognized alternatively as mediators of neurodegeneration^24–35^ or as sentinels protecting the brain from neurodegenerative pathogenesis^25,36^. Studies on the role of T cells in neurodegeneration is largely limited to proteinopathies like amyloid β, hyperphosphorylated tau, or α-synuclein deposition, and most of these studies have been conducted at juvenile or young ages in mice. Several single-cell RNA sequencing (scRNA-seq) studies have been undertaken to capture the cellular heterogeneity within the mouse brain and in health and disease.^37^ One of the largest, the mouse brain cell atlas generated by the Allen Brain Institute includes transcriptomic profiling of more than 4 million individual cells; however, only about 300 cells are T cells and all mice were between 7-10 weeks of age.^26^ Some scRNA-seq studies have investigated aging mouse brain and have exceptional resolution for neurons, microglia, astrocytes, and oligodendrocytes but contain almost no T cells.^32,34,38^ The lack of T cells in these large scRNA-seq studies in mice and human brains is not reflective of a true lack of T cells within the brain because imaging and flow cytometry studies have demonstrated that T cells within the human brain may outnumber macrophages.^39,40^ The lack of high-dimensional data on T cells and their interaction with microglia in the aging brain represents a critical gap in knowledge, especially given the rapid pace of therapeutic development of T cell-targeting drugs for autoimmune disease and cancer which could be repurposed for neurodegenerative disease.

In summary, addressing neurodegeneration and the contributions of immunity to brain health requires a comprehensive, multi-modal examination of all immune cells in the brain, including T cells, within an age-appropriate context reflective of the period of greatest neurodegenerative disease risk in humans. To accomplish this, we used a multi-modal approach including high-dimensional multiplex imaging, spectral flow cytometry, and Split Pool Ligation-based Transcriptome sequencing (SPLiT-seq)^41^, to generate a comprehensive dataset encompassing microglia, myeloid cells, and lymphoid cells in both the young and aged brain in health and a model of VaD. We compared and confirmed key microglial phenotypic shifts from murine SPLiT-seq data with human microglia in health and VaD. Notably, microglial ligand-receptor analysis with T cells identified antigen-presentation and co-stimulatory transcriptional profiles linked to a novel subset of granzyme k-enriched cytotoxic CD8^+^ T cells via CD28 signaling. These findings support a model where microglia-to-T cell interactions drive inflammation in the aging brain in VaD.

## RESULTS

### Immune Cells Make Up a Significant Component of the Brain in Age and VCID

Understanding how resident immune cells influence peripheral immune cells in the brain involves complex interactions between diverse cell types.^13^ Most chronic diseases manifest in old age, and aging itself significantly alters both resident and peripheral immune cells in the brain. A complicating factor is that peripheral immune cells like macrophages and T cells are significantly outnumbered in the brain by neurons.^42^ Thus, most scRNA-seq atlases in the murine brain contain very few peripheral immune cells, which severely limits our understanding of how peripheral immune cells in the brain change by age and in disease. To understand how age modifies both the resident and peripheral immune landscape in the brain during health and in disease, we collected brains from mice. Mice were classified as either young (3-months) or old (21-months), at which time half of each were subjected to either bilateral carotid artery stenosis (BCAS) surgery^43^ or sham surgery then maintained for another 30-days.

We used the BCAS surgery to model cerebrovascular inflammation common in VaD and VCID.^5^ Briefly, we placed microcoils (0.17 mm inner diameter, 2–3 mm length) on each common carotid artery for 30 days. Thus, brains were harvested from mice from four groups: 4-month control, 4-month VaD, 22-month control, and 22-month VaD, with 18 mice per group. After 30 days, animals were perfused, and whole brains were harvested. Brains from mice in each group were harvested for 26-plex multi-parameter immunofluorescent imaging, high dimensional spectral flow cytometry including 36 antibody targets, and split-pool ligation transcriptomic single-cell RNA-sequencing (SPLiT-seq)^41^ (Figure 1A). For the SPLiT-seq samples, we homogenized and digested the brain, immunolabeled for live, singlet immune cells (CD45) and T cells (CD3e). We sorted and collected all CD45^hi^CD3^+^ T cells in addition to CD45^hi^CD3^−^ peripheral immune cells and CD45^int^ microglia (resident immune) cells (Figure S1). We also confirmed that CD3^+^ T cells expressed canonical CD4 and CD8 surface receptors for helper and cytotoxic T cells, respectively (Figure S1). Sorted cells were then fixed in batches prior to library preparation for SPLiT-seq^41^ and subsequent data analysis (Figure 1A).

**Figure 1.**
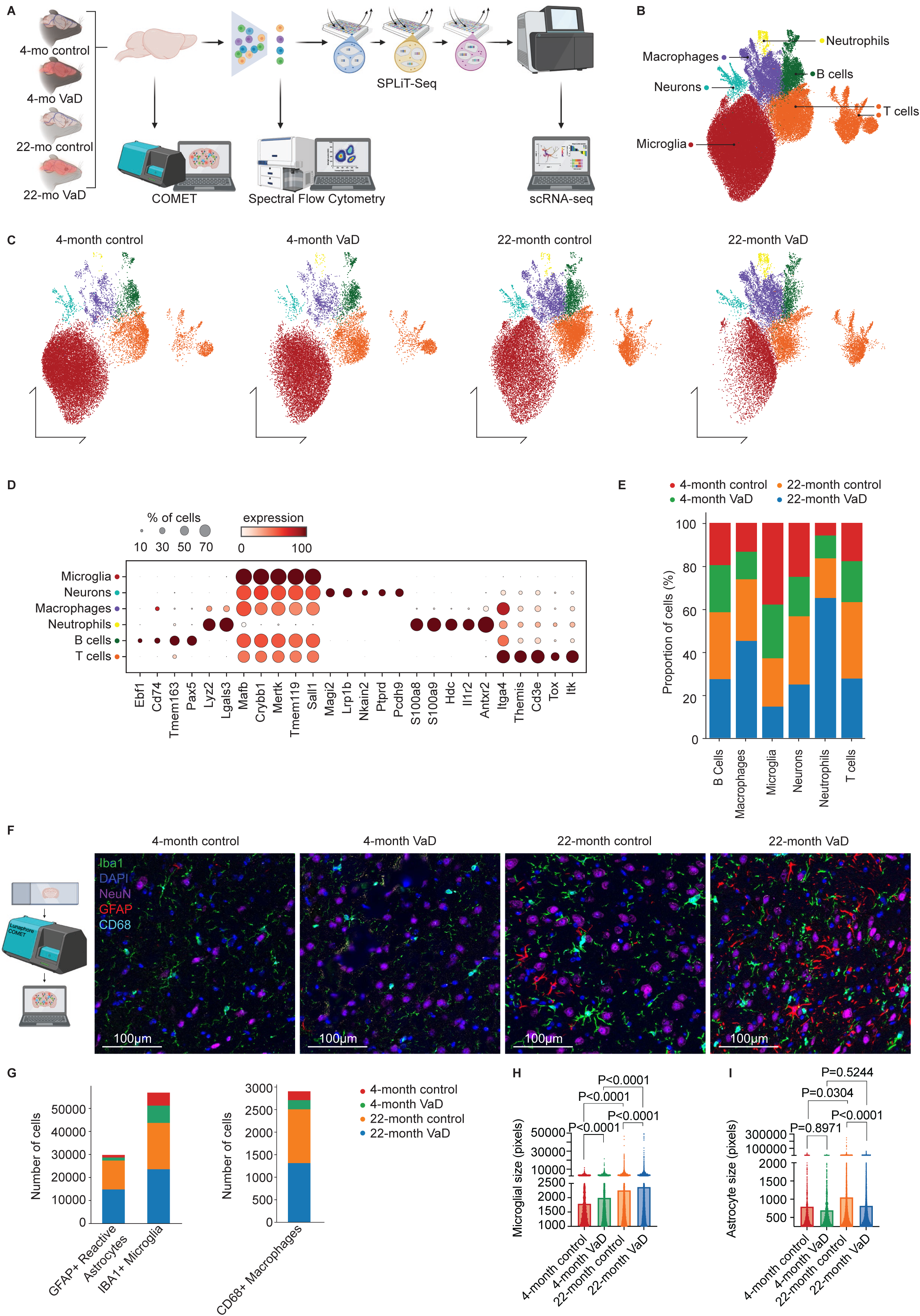
Single-Cell Immune Atlas of the Aged Mouse Brain Reveals Dynamic Changes Across Experimental Groups. (A) Experimental design: 52 female C57BL/6N wild-type mice (N=13 per group) aged 4 months (young) or 22 months were grouped into 4-month Control, 4-month VaD (vascular dementia), 22-month Control, and 22-month VaD, N=13/group. Whole mouse brain was used for preparing samples for Lunaphore COMET multiplex imaging, spectral flow cytometry, and split pool ligation transcriptomics (SPLiT-seq). (B) Unsupervised Leiden clustering and annotation revealed 6 metaclusters. (C) Cell metaclusters are shown by age and treatment groups. (D) Dot plot displays marker gene expression for annotated cell types of each metacluster. (E) Stacked bar plot showing the relative distribution (%) of major cell types across experimental groups. Each color represents a distinct experimental group. Cell proportions were calculated relative to the total number of cells within each identified immune cell type. (F) Representative multiplex immunofluorescent images from Lunaphore COMET imaging for each group in the murine brain striatum demonstrating staining for nuclei (DAPI), microglia (Iba1), neurons (NeuN), astrocytes (GFAP), and macrophages (CD68). Scale bars = 100µm. (G) Stacked bar plot of total number of reactive astrocytes, microglia, and macrophages from COMET imaging within each group. (H and I) Quantification of the size of microglia (H) and astrocytes (I) from COMET imaging for each group. N=3 biological replicates with 4-5 brains pooled per biological replicate for each group (total=52 brains). n=69,614 cells total.

We used Leiden^44^ clustering with 70,000 total immune cells, and we identified 6 unique metaclusters (Figure 1B). Here, we refer to brains that underwent BCAS surgery as “VaD”; however, “VCID” could also be appropriate. There were differential proportions of cells evident among the unique metaclusters (Figure 1B) which showed unique representation by age and treatment (Figure 1C). Using marker gene expression for each of the metaclusters, we identified microglia (*Tmem119*^+^), T cells (*Cd3e*^+^), macrophages (*Mertk*^+^), B cells (*Pax5*^+^ and *Tmem163*^+^), neutrophils (*S100a8*^+^ and *S100a9*^+^), and neurons (*Lrp1b*^+^ and *Pcdh9*^+^) (Figure 1D). About 60% of the cells were identified as microglia which were more prominent in young brains (Figure 1E). Of the remaining cells 25% were identified as T cells, 5% were B cells, macrophages, and neutrophils, which were all more abundant in aged brains (Figure 1E). The number of microglia declined with VaD in both young animals (∼60% in control and ∼55% in VaD) and aged animals (50% in control and 40% in VaD; Figure 1E). Myeloid cells were more enriched in aged VaD brains (Figure 1E). These findings highlight the important role that aging plays on polarization and recruitment of subsets of peripheral and resident immune cells in the brain; however, the total abundance of cell types is dependent on digestion and enrichment of immune cells from the brain prior to SPLiT-seq.

To gain a better understanding of abundance and representation of microglia, macrophages, and astrocytes, which were not captured in our SPLiT-seq analysis, we performed multiplex immunofluorescent imaging using Lunaphore COMET^TM^. We stained tissue sections and performed COMET imaging for major immune cells including microglia, astrocytes, and macrophages, in addition to neurons, nuclei, and phenotypic markers (Figure 1F). We found that GFAP^+^ cells (reactive astrocytes), Iba1^+^ cells (microglia), and CD68^+^ (macrophages) were all more enriched in the aged brain (Figure 1G). We further examined the size of individual microglia (Figure 1H) and astrocytes (Figure 1I) by group and age and found that microglial size significantly increases with age and treatment while astrocyte size increases with age but decreases with VaD in the aged brain. These data demonstrate significant reactivity of both resident microglia and peripheral immune cells within the aged brain with subtler inflammatory alterations due to treatment.

### Aging Dramatically Alters Immune Cell Heterogeneity in the VCID Brain

We next analyzed the cellular subset composition and heterogeneity for the major cell types identified in our SPLiT-seq dataset. Microglia (36,900 cells) included quiescent cells that were classified as homeostatic, characterized by higher expression of *Tmem119*, *Sall1*, *P2ry13*, and heat shock genes (Figures 2A&B).^45^ Homeostatic microglia were classified into four clusters including two that are more classically homeostatic in transcriptomic profile (0_Homeostatic 1 and 6_Homeostatic 2) with the highest *Sall1* and heat shock gene expression. We also identified a homeostatic microglial cluster with high expression of heat shock and DNA repair genes (*Hspa1a*, *Hsp90aa1*, *Dnajb1*), along with canonical homeostatic markers such as *Sall1* and *Fcrls* (Figure 2B) which we termed 2_Stress-response.^46^ The final homeostatic cluster (Inflammasome-related Microglia) specifically showed elevated expression of *Nlrp3*, *Casp4*, and *Stat3*, reflecting involvement in innate immune and inflammasome pathways (Figure 2B).^47^

**Figure 2.**
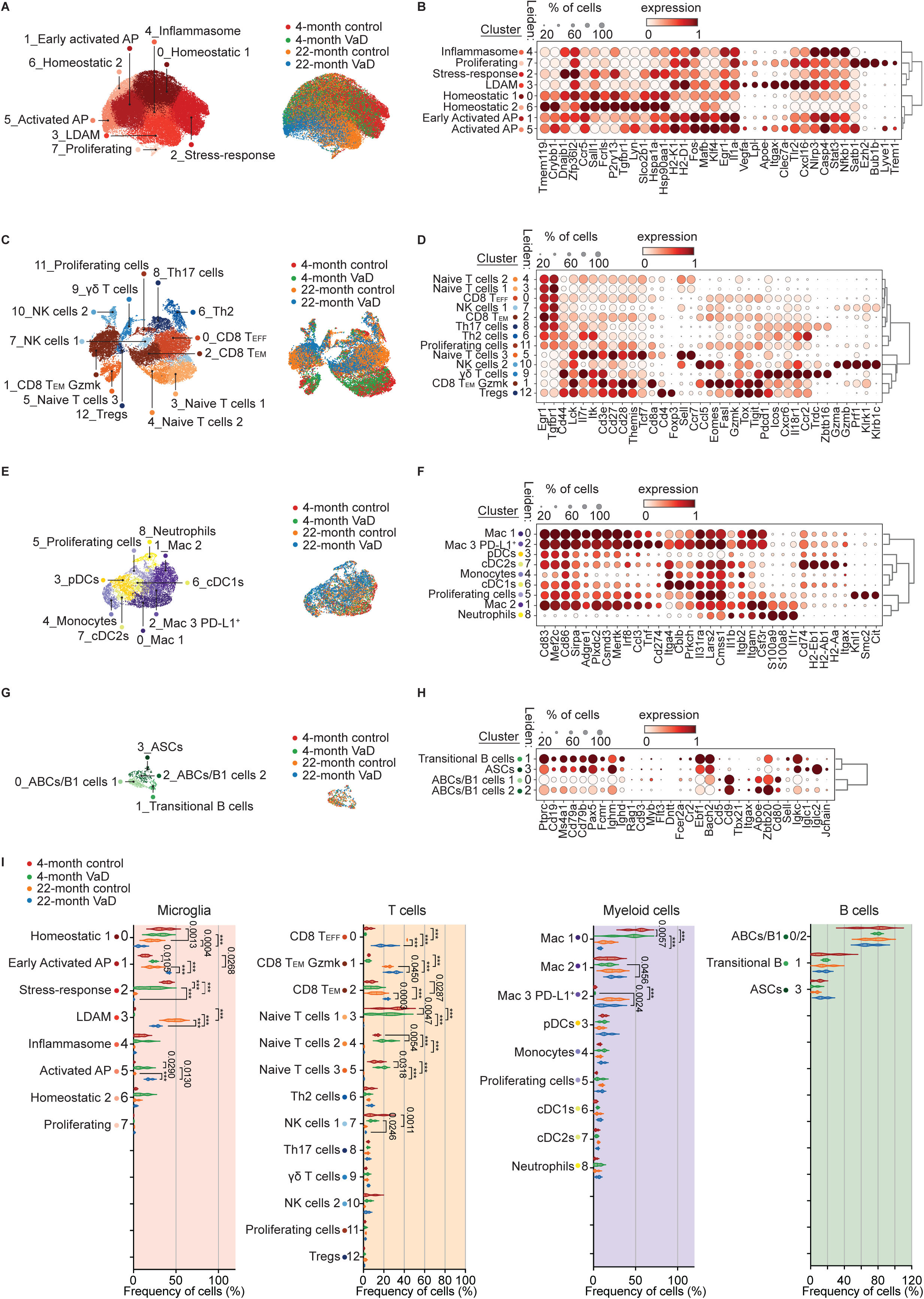
Brain Immune Cell Classification Reveals Heterogeneity by Age and Vascular Dementia. Cells from SPLiT-seq from 52 female C57BL/6N wild-type mice (N=13 per group) aged 4 months (young) or 22 months were grouped into 4-month Control, 4-month VaD (vascular dementia), 22-month Control, and 22-month VaD, N=13/group were subset into metaclusters of microglia (A), T cells (B), myeloid cells (C), and B cells (D). Each metacluster was sub-clustered and cell subsets annotated and shown as UMAP embedding on left (A, C, E, G) and by age and treatment group in middle (A, C, E, G). Dot-heatmaps for each sub-cluster demonstrate averaged gene expression and percentage positive gene expression along with K-means dendrogram clustering of sub-clusters on right (B, D, F, H). (I) Violin plots of immune subset composition for microglia, T cells, myeloid cells, and B cells across four experimental groups. Violin plots represent mean ± quartiles. Statistical significance among groups were assessed by two-way ANOVA with Fishers Least Significant Difference post-hoc test with p-values indicated above each relevant comparison. N=3 biological replicates with 4-5 brains pooled per biological replicate for each group (total=52 brains). n=69,614 cells total. n=36,900 microglia, n=16,479 T cells, n=7,913 myeloid cells, and n=631 B cells.

Next, we examined expression of common pro-inflammatory genes *Apoe*, *Cybb*, *Axl*, *Il1b*, and *Tnf*. We found three clusters with reduced *Tmem119* relative to the homeostatic clusters (0, 2, 4, and 6) suggesting these 3 subpopulations undergo a shift away from a homeostatic state. We found one cluster, Early Activated AP Microglia, had moderate upregulation of inflammatory genes *Il1a*, *Nlrp3*, and *Nfkb1*, and major histocompatibility complex (MHC)-I genes *H2-K1* and *H2*-*D1*, but retained some homeostatic markers (*Sall1*, *Hspa1a*, *Hsp90aa1*, *Dnajb1*), suggesting a partially activated state (Figures 2A&B). The other two pro-inflammatory clusters were characterized by even further diminished expression of homeostatic genes linked to DNA repair/heat shock proteins, along with higher levels of inflammatory genes such as *Trem1*, *Il1a*, *Apoe*, *Clec7a*, *Nlrp3*, *Nfkb1*, *Egr1*, and *Tlr2*, and MHC-I genes *H2-K1* and *H2*-*D1* consistent with a disease-associated and/or inflammatory phenotype (Figure 2B). We annotated these as 5_Activated Antigen Presenting (AP) microglia and 3_Lipid Droplet Associated Microglia (LDAM) based on differential expression of lipid handling genes such as enrichment of *Lpl* in the LDAM cluster (Figures 2A&B). We annotated cluster 3 as LDAM instead of disease-associated microglia because this cluster exists within all groups including young and aged control mice (Figure 2B). Although LDAM and disease-associated microglia are semantic labels for similar microglial phenotypes, we used LDAM due to the lack of true disease in aged control mice here.^12,48,49^ The final microglial cluster, 7, was the smallest cluster which is enriched for genes related to the cell-cycle and ribosomal biogenesis which are indicative of proliferating cells (Figure 2B).

Given the important role of T cell–microglia interactions in neuroinflammation and aging, we next focused on T cells, which represented the second most abundant immune cell population in our dataset. We identified 13 distinct T cell clusters, primarily within the CD4 and CD8 αβ T cell lineages with smaller clusters of T cells in the γδ T cell lineage and natural killer (NK) lineage (Figure 2C). Naïve T cells are un-differentiated surveillance T cells that can recognize a wide variety of antigens by antigen-presenting cells; however, naïve T cell numbers decline as a function of age due to thymic involution in both mice and humans.^50–52^ To determine which clusters in the brain were naïve T cells, we examined expression of *Ccr7* and *Cd44*. *Ccr7* is a hallmark naïve T cell marker and naïve T cells should not express *Cd44* which is indicative of antigen experience (Figure 2D). Memory T cell subsets were determined based on loss of *Ccr7* expression and gaining expression of *Cd44*.

We found three clusters of naïve T cells characterized by expression of *Ccr7* and *Sell* (encoding CD62L) and lacking expression of antigen experience markers like *Cd44* (Figures 3C&D).^50^ These were αβ T cells of both CD4 and CD8 lineages and were almost exclusively restricted to young brains (Figure 2C). In the aged brains, we identified distinct clusters of effector and memory CD8^+^ T cells with many expressing *Gzmk* (clusters 0, 1, and 2; Figures 3C&D). *Gzmk* encodes for granzyme K which is a tryptase-class serine protease localized in the cytotoxic granules of CD8⁺ T cells and, in humans some invariant T cell sub-types like mucosal associated invariant T cells (MAITs).^53–56^ It contributes to non-canonical cytotoxic pathways by inducing target cell apoptosis and promoting pro-inflammatory signaling through substrate cleavage.^55–59^ Granzyme K-expressing T cells were recently identified to accumulate in several tissues in mice and human with age.^56,57,60–62^ We annotated the effector and memory CD8^+^T cells as CD8^+^ T effector (T_EFF_), CD8^+^ effector memory (T_EM_) cells, and *Gzmk*-expressing CD8^+^ T_EM_ cells (T_EM_ Gzmk) based on differential expression of *Gzmk* and terminal expression markers including chemokine receptors (Figure 2D). The CD8^+^ T_EM_ Gzmk cells exhibited high expression of genes suggesting antigen experience and exhaustion^63^ including *Cd44*, *Lag3*, *Tox*, and *Pdcd1* along with inflammation and cytotoxicity including *Ccl5*, *Fasl*, *Eomes*, *Tigit*, *Lck*, and *Icos* (Figure 3F). We also identified two clusters of natural killer cells (NK cells), a γδ T cell cluster, a cluster of CD4^+^ regulatory T cells (Tregs), CD4^+^ T cell clusters for both Th2 and Th17 lineage, and proliferating T cells.^64^ These were identified based on expression of γδ T cell receptor (TCR) expression, *Cd4*, *Cd8a*, *Cd8b1*, and genes associated with invariant T cells such as *Zbtb16*, *Klrk1*, *Klrb1c*, and *Il18r1* (Figure 2D).

**Figure 3.**
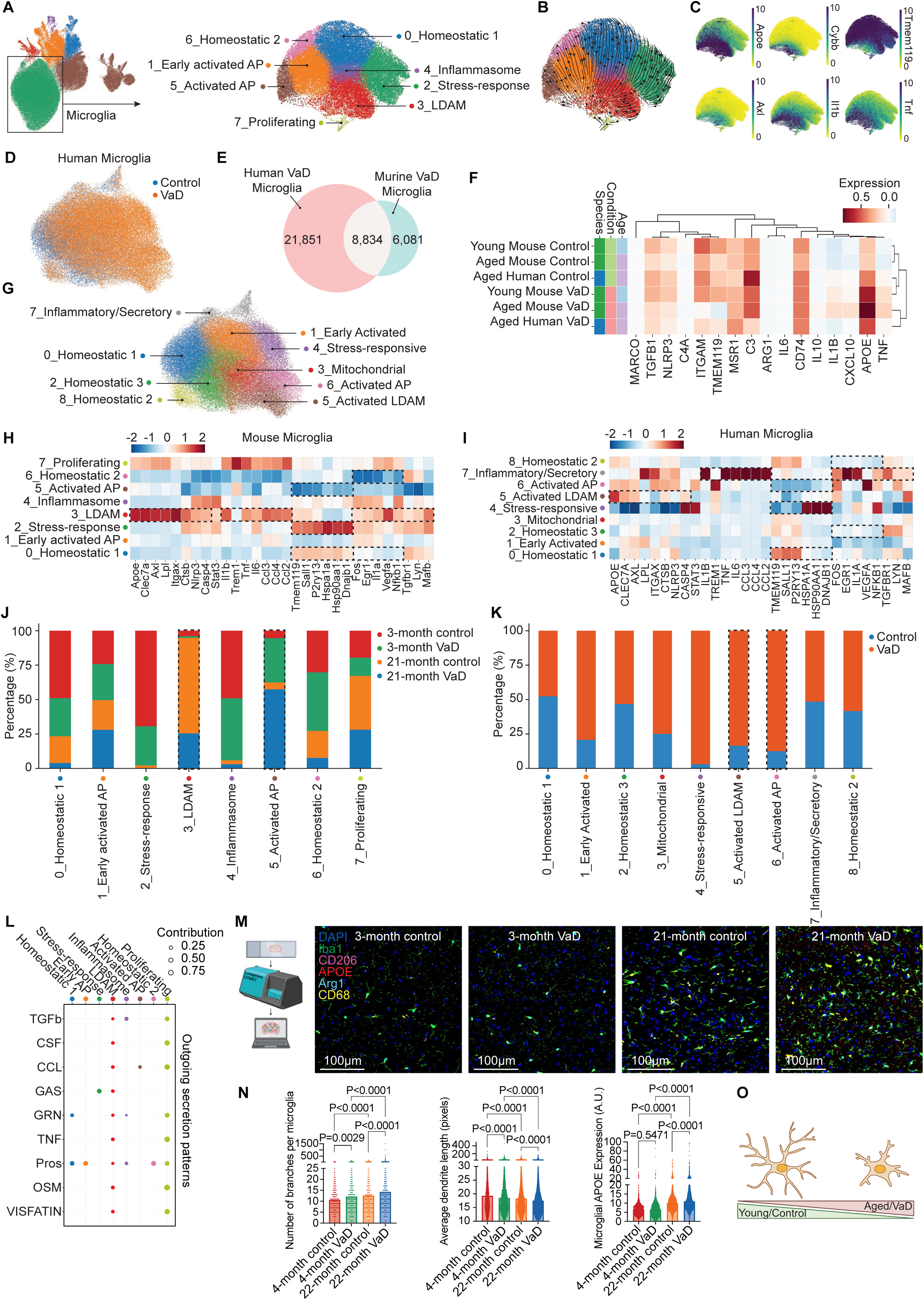
Dynamic Microglial Heterogeneity and Transcriptional Trajectories in Vascular Dementia. (A) Unsupervised Leiden sub clustering of microglia populations resolved 8 transcriptionally distinct immune cell clusters shown by UMAP embedding. (B) RNA velocity streamlines projected onto UMAP embeddings, with arrows indicating transcriptional trajectories and colors reflecting Leiden clusters. (C) UMAP embeddings showing gene expression of key homeostatic and inflammatory classification genes (*Apoe*, *Cybb*, *Tmem119*, *Axl*, *Il1b*, *Tnf*). (D) Human microglial single-cell RNA-seq analysis from healthy brain and vascular dementia (NCBI: GSE213897) shown by group via UMAP embedding. (E) The number of overlapping and non-overlapping genes from the human microglial dataset and our SPLiT-seq murine microglial dataset by Venn diagram. (F) Heatmap of key microglial sub-classification gene expression organized by K-means clustering for human and murine microglial genes together. (G) Human microglial classification by marker gene expression plotted as UMAP embedding. Heatmap of gene expression of mouse (H) and human (I) microglia by sub-cluster. Stacked bar plot of mouse (J) and human (K) microglia by treatment group for each microglial sub-cluster. Relative proportions (%) of the total cell count per cell type are shown. (L) Outgoing secretion patterns of murine microglial SPLiT-seq sub-clusters with strength of contribution. (M) Representative multiplex immunofluorescent images of microglia from Lunaphore COMET imaging for each group in the murine brain striatum demonstrating staining for nuclei (DAPI), microglia (Iba1) and microglial phenotype markers: CD206, APOE, Arg1, and CD68. Scale bars = 100µm. (N) Quantification of the number of dendrite branches of microglia, average length of dendrite branch per microglia, and average APOE expression per microglia by murine treatment group. Statistical significance determined by one-way ANOVA with Sidak post-hoc test. (O) Schematic of murine microglial morphology change by age and treatment in mice. N=3 biological replicates with 4-5 brains pooled per biological replicate for each group (total=52 brains). n=36,900 microglial cells.

Brain-resident macrophages have been shown to present antigen to CD4^+^ T cells in Parkinson’s disease models.^65,66^ Depletion of microglia and brain-resident macrophages enhances phospho-tau deposition in P301S^−/-^ models leading to premature death compared with controls.^36,67^ Brain-resident macrophages also change considerably with age and those changes could have impacts on VCID onset and progression.^13,68^ Thus, we examined the 7,913 non-microglial myeloid cells in our dataset. We identified 9 distinct clusters of myeloid-lineage cells (Figure 2E). We examined the marker genes for each of the clusters of cells and identified 3 clusters of macrophages, 3 clusters of dendritic cells, a cluster of monocytes, a cluster of neutrophils, and a cluster of proliferating cells (Figure 2F). Cluster identities were confirmed by expression of common marker genes like *S100a8* for neutrophils, *Mertk* for macrophages, *Kif15* for proliferating cells, and *Itgam* (encoding CD11b) for myeloid cells (Figure 2F). We noted that cluster 2, which were identified as macrophages had high expression of *Cd274* which encodes for PD-L1 compared to the other clusters (Figure 2F).

B cells have been identified in the brain in an Alzheimer’s disease mouse model^69,70^; however, less is known about their composition in aging or VaD.^71^ Initial sub-clustering of B cells revealed some other cell types within the B cell metacluster. We used stringent criteria to isolate true B cells present in the brain of aged and young mice. Only cells expressing constituents of the BCR (*Igh* and *Igl* genes) as well as BCR co-receptors (*Cd79a* or *Cd79b*) were included for downstream analysis, additionally any cells expressing canonical T cell markers (*Cd3e*, *Cd4*, or *Cd8*) were removed from downstream analysis. We identified 631 remaining B cells (Figure 2G) with 4 transcriptionally distinct subpopulations of B cells present. All clusters exhibited expression of canonical B cell genes including *Cd19*, *Ms4a1*, *Cd79a*, and *Cd79b*, indicating the purity of our population (Figure 2H). Genes regulating the B cell program such as *Pax5*, *Bach2*, and *Ebf1* were highly expressed across clusters supporting the idea that the majority of B cells present are not terminally differentiated (Figure 2H).^72^ Cluster 3 exhibited the high expression of the BCR genes *Ighm*, *Iglc1*, *Iglc2*, and *Igkc*, which suggests these are antibody secreting cells (ASCs).^72^ Cluster 1 was characterized by mature B cell genes^73,74^, were negative for key regulators of B cell development including *Rag1*, *Cd93*, *Myb*, *Flt3*, and *Dntt*, and only a fraction expressed *Cr2* and *Fcer2a*^75^ which suggests these are transitional B cells.^73,74^ B cell clusters 0 and 2 had enrichment of several genes associated with age-associated B cells (ABCs) in the brain including *Zbtb20*, *Cd80*, and *Apoe*.^76^ These cells also exhibit high expression of *Cd9*, a marker gene for innate-like B1 cells^77^ which have not been extensively described in the brain but have been observed in the meninges of mice (Figures 2G&H).^76^ This analysis demonstrates the heterogeneity of transcriptionally distinct B cell populations in the brain.

We next sought to determine whether the sub-clusters of either microglia, T cells, myeloid cells, or B cells were statistically significantly altered by age or treatment. We found that among microglia, the Homeostatic_1 and Stress-responsive clusters were significantly reduced by age in both treatment groups (Figure 2I). We found that the VaD treatment also further reduced the number of microglia in these clusters (Figure 2I). The Early Activated AP and LDAM clusters were both significantly enriched in aged brains compared to young with Early Activated AP being more enriched in Aged VaD while LDAMs were more enriched in aged control brains compared to young treatment controls (Figure 2I). The more terminal microglial cluster, Activated AP cells, were uniquely enriched in the VaD treatment group compared to controls in young brains and even further enriched in the Aged VaD group, suggesting a unique VaD-related signature of accumulation, independent of age (Figure 2I).

T cell accumulation in the brain showed significant shifts by both age and treatment as well. All 3 Naïve T cell clusters, along with NK cells_1 were enriched in young brains with almost no naïve T cells found in the either aged group (Figure 2I). The 3 activated clusters of CD8^+^ T cells were enriched in the aged brains in both treatment groups compared to young with the 2 CD8^+^ T_EM_ clusters (CD8 T_EM_ Gzmk and CD8 T_EM_) being more enriched in the aged VaD group compared to the aged control group (Figure 2I). These results suggest that both age and VaD treatment skew memory CD8^+^ T cell accumulation in the brain.

Among the myeloid clusters, the most abundant sub-cluster, Macrophage_1, was enriched most in young control brains followed by Young VaD brains compared to both aged groups suggesting an aging-specific accumulation (Figure 2I). The other 2 macrophages sub-clusters, Macrophage_2 and Macrophage_3 PD-L1^+^, were enriched in Aged VaD brains compared to Young VaD brains (Figure 2I). The PD-L1^+^ Macrophage_3 cluster was also enriched in the aged control brains compared to young control suggesting that this cluster is influenced primarily by age rather than treatment (Figure 2I). The rest of the myeloid clusters and the B cell clusters were very small clusters and were not statistically significantly altered by age or treatment.

### Microglial Reveal Age and Disease-Driven Inflammatory Shifts

In the setting of tauopathy, amyloid-β deposition, and aging, microglia tend to shift from homeostatic or resting phenotype to disease-associated microglial or lipid droplet-associated microglial (LDAM) phenotype.^12,17,48,78^ To determine how microglia are impacted by VaD, which occurs mostly in old age, we isolated the microglial metacluster to better understand how microglia transition between different cell states over time (Figure 3A). We performed RNA velocity analysis^79^ which positioned the homeostatic clusters (6, 0, and 2) at the “origin,” serving as an entry point for transcriptomic velocity flow (Figure 3B). In contrast, LDAMs, Early Activated AP Microglia, and Activated AP Microglia (clusters 3, 1, and 5 respectively) appeared as the most differentiated or “terminal” states (Figure 3B). These observations are consistent with a model in which homeostatic microglia can progressively transition to activated or disease-associated phenotypes, especially under conditions of aging or vascular risk.^80,81^ We further examined specific anti- and pro-inflammatory gene expression known to accumulate in either homeostatic or activated microglia. The canonical anti-inflammatory microglial gene, *Tmem119*, was more widely expressed in the homeostatic clusters whereas pro-inflammatory genes like *Apoe*, *Cybb*, *Axl*, *Tnf*, and *Il1b* were more highly expressed in the activated microglial clusters (Figure 3C). In support of this, the least differentiated clusters were enriched in young brains, whereas the more differentiated proinflammatory subsets predominated in aged mice (Figure 2I). Thus, our findings suggest that homeostatic or quiescent microglia become progressively activated and accumulate with age, contributing to the loss of protective homeostatic microglia in older brains.

To validate our scRNA-seq results at the protein level, we performed spectral flow cytometry of microglia isolated from a contemporaneous cohort of female C57BL/6N mice. We examined four groups (4-month-old control, 4-month-old VaD, 22-month-old control, and 22-month-old VaD), with n=5 mice in each control group and n=4 mice in each VaD group. Perfused, whole brains were harvested, and immune cells were enriched and stained using a panel of antibodies targeting both homeostatic and proinflammatory microglia (Figure S2A). We gated live CD45^int^ CD11b^+^ cells to identify microglia, then performed PCA, Leiden clustering, and UMAP to identify microglial sub-populations.^82,83^ We identified eight distinct clusters of microglia (Figure S2B) based on surface protein expression (Figure S2C). The major homeostatic clusters (clusters 0, 1, 3, 4, and 5) were more abundant in younger animals and were defined by lower expression of chemokine markers CXCR3, CCR2, CCR5 as well as less Siglec H and CD64 (Figure S2D). The intermediate and proinflammatory clusters (clusters 2, 6, and 7) were greater in aged mice (Figure S2D), and all microglia were defined by intermediate CD45 expression, low CD11b expression, and high CX3CR1 expression compared with peripheral immune cells like T cells and macrophages. Thus, the spectral flow cytometry data corroborate our transcriptomic results, demonstrating a shift toward proinflammatory microglia accumulation in older brains and in VaD relative to young controls.

### Microglia in aged mouse brain share features of aged human brain in VaD

In order to compare our findings to human disease, we examined a comprehensive public scRNA-seq dataset of human brain cells in disease-free and VaD patients.^84^ This dataset included five controls (mean age 82.6 years) and five individuals with VaD (mean age 80.2 years). There were no significant differences in age between the groups.^84^ When we compared genes expressed in the human brain microglia and murine brain microglia, we found 8,834 corresponding genes that overlapped between the two datasets (Figures 3D&E). We examined key marker genes and inflammatory genes across the four murine groups and two human groups and performed K-means clustering to determine group similarities. This analysis grouped the aged mouse VaD group with human VaD group as well as the young mouse control group with the aged mouse control groups in an unbiased manner (Figure 3F). We next identified marker genes for the eight murine microglial sub-clusters from our dataset that had corresponding genes to the nine human microglial sub-clusters that we identified after performing Leiden clustering on the human microglial dataset and annotated clusters based on similar marker gene expression (Figures 3F&G). We identified similar clusters across the murine and human microglial sub-clusters. We found that the murine LDAM cluster was similar in expression to two of the human microglial sub-clusters which we identified as Activated LDAM and Inflammatory/Secretory clusters based on marker gene expression (Figures 3H&I). We also found similarities in the quiescent homeostatic microglial clusters between the murine and human microglial datasets. When we examined these clusters by group (mice and humans) and age (mice only), we found that the similar accumulation patterns persisted for some of the microglial sub-clusters. The two pro-inflammatory murine microglial sub-clusters, LDAMs and Activated AP microglia, were enriched in aged mice and VaD mice, respectively (Figure 3J).

Similarly, the pro-inflammatory and activated clusters identified in the human microglial sub-clusters, Activated LDAMs and Activated AP microglia, were similarly enriched in the human VaD group (Figure 3K). Interestingly the murine quiescent sub-cluster, stress-responsive microglia, which was enriched in young control mice corresponded to a sub-cluster with a similar transcriptomic profile in the human microglial dataset; however, these were highly enriched in the human VaD group (Figures 3J&K). There were notable differences between the two stress-responsive microglial sub-clusters including high *Tmem119*, *Sall*, and *P2ry13* which were particularly low in the human dataset (Figures 3H&I). This could indicate that the human stress-responsive microglial sub-cluster is in a more transitionally activated state than the corresponding murine cells. Nevertheless, these data indicate strong concordance between murine microglia and human microglia in the setting of VaD, particularly when at older ages in both mice and humans.

### Age and VaD Influence Microglial Intercellular Communication and Exhibit Distinct Pro-Inflammatory Profiles

To gain a deeper understanding of microglial changes in age and VaD, we used the CellChat package on our murine dataset, which infers cell-cell communication networks by identifying and quantifying potential ligand-receptor interactions based on gene expression profiles.^85^ We found that LDAMs and proliferating microglia displayed the highest outgoing signals via multiple ligands predicted to act on other clusters including TGF, CSF, CCL, and TNF pathways (Figure 3L). Given our transcriptomic findings that microglia in aged brains may be more secretory and prior reports demonstrating microglial dendrite rarefaction and soma expansion during activation and cytokine secretion, we examined microglial morphology in our COMET imaging data (Figure 3M). Along with increased size (Figure 1H), we found that the number of dendrite branches per microglia increases with age and treatment while the average length of dendrites decreases (Figure 3N). At the same time, protein expression of APOE, a strong indicator of microglial activation and phagocytic function is greater in old age and the setting of VaD, particularly at old age (Figure 3N). Together, our multimodal data analysis indicates that with age and VaD, quiescent/homeostatic microglia are lost in favor of more secretory and phagocytic microglia (Figure 3O).

Within all four murine groups, we identified microglia associated with each transcriptomic cluster within our multiplex imaging data (Figure 4A). Microglia with more and longer dendrites were characterized by less protein expression of APOE and CD68 but greater Arg1 and CD206. In contrast, microglia with larger soma and shorter dendrites were characterized by greater APOE and CD68 expression but less Arg1 and CD206 (Figure 4A). These representative images corroborate our transcriptomic findings and further demonstrate that microglial morphologic heterogeneity is common within young and aged mice in health and in VaD. These data suggest that microglia may be particularly plastic with respect to phenotype and morphology and thus represent promising targets therapeutically for neurodegenerative disease.

**Figure 4:**
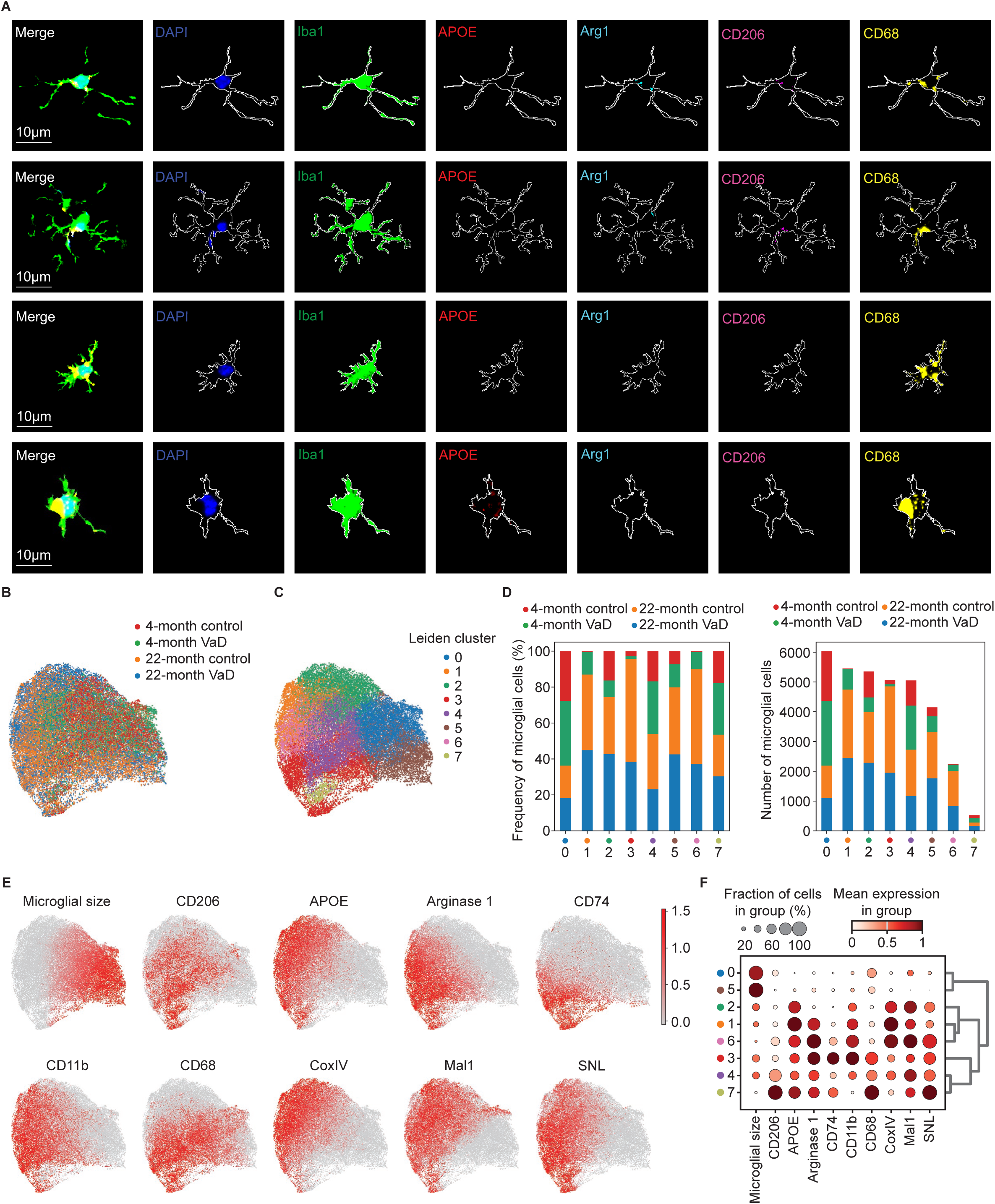
Microglial Morphology is Heterogeneous in Aging and Vascular Dementia. (A) Representative multiplex immunofluorescent images of microglia from Lunaphore COMET imaging in the murine brain striatum demonstrating staining for nuclei (DAPI), microglia (Iba1) and microglial phenotype markers: CD206, APOE, Arg1, and CD68. Scale bars = 100µm. Microglia COMET imaging protein expression were quantified and displayed as UMAP embedding (B) and clustered by unsupervised Leiden algorithm (C). (D) Stacked bar plot of mouse microglia by age and treatment demonstrating frequency (%) of microglia by cluster and total number of each cluster. (E) UMAP embedding of microglial protein expression from COMET imaging for microglial phenotype. (F) Dot-heatmap for each sub-cluster of microglial demonstrates averaged protein expression and percentage positive protein expression along with K-means dendrogram clustering of sub-clusters on right. N=1 biological replicate per group (total=4 brains). n=33,882 microglia.

We further examined microglial size along with 9 other phenotypic markers from our multiplex imaging data and performed un-supervised Leiden clustering which identified 8 distinct clusters based on imaging data (Figures 4B-D). The clusters were defined by specific marker genes with clusters enriched in known pro-inflammatory associated proteins like APOE to be more associated with aged and VaD samples in contrast to the more anti-inflammatory markers like Arg1 (Figure 4E&F). Interestingly, we identified that some clusters were enriched in expression of the mitochondrial protein COXIV, which is indicative of mitochondrial content. These clusters (1, 2, and 6) were particularly more abundant in the aged brain images suggesting these microglia may be more metabolically active (Figures 4D&F). This is consistent with the microglial metabolic requirements associated with cytokine secretion and phagocytic activity.

### Aged Brains Exhibit Enrichment of Terminally Differentiated Pro-inflammatory *Gzmk*-Expressing CD8^+^ T Cells

Given the abundance of T cells within the brain, we further examined the T cell sub-clusters (Figure 5A). We noted that naïve T cells were only appreciable in young brains but were nearly absent in aged brains (Figure 5B). The aged brains primarily exhibited memory CD8^+^ T cells (clusters 0, 1, and 2; Figure 5B). We found that aged brains in both VaD and control groups possessed large numbers of *Gzmk*^+^ effector memory CD8^+^ T cells, cluster 1: CD8 T_EM_ Gzmk (Figure 5B). We confirmed our finding of greater numbers of memory (CD44^+^) CD8^+^ T cells in both aged mouse brains compared with young brains in our COMET dataset (Figure 5C).

**Figure 5:**
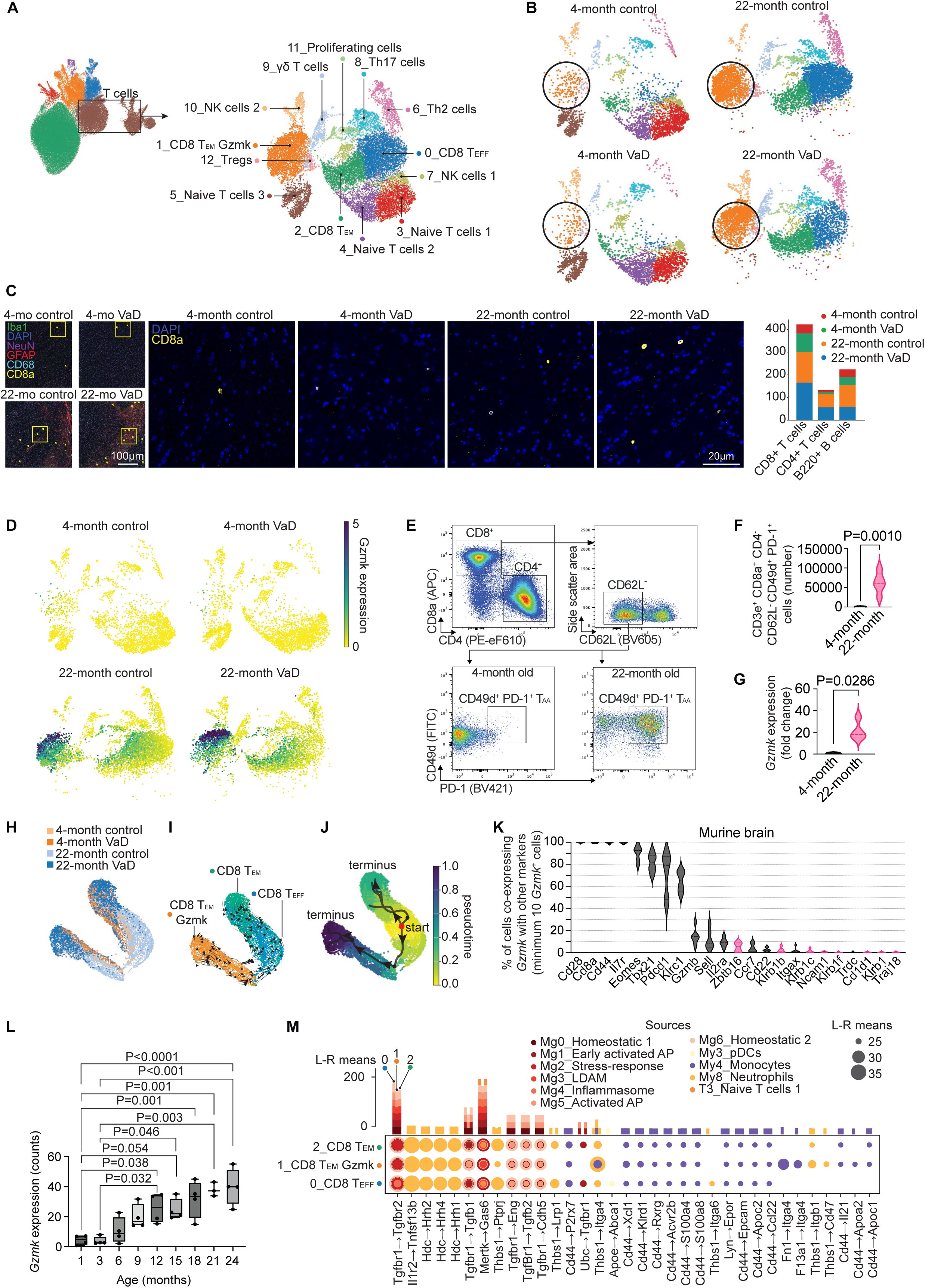
Accumulation of *Gzmk*-expressing CD8^+^ T Cells in the Aging Brain. (A) Unsupervised Leiden sub-clustering of T cell populations resolved 13 transcriptionally distinct clusters and plotted by UMAP embedding. (B) The distribution of Leiden clusters by age and VaD status plotted by UMAP embedding. (C) Representative multiplex immunofluorescent images of CD8^+^ T cells from Lunaphore COMET imaging for each group in the murine brain striatum demonstrating staining for nuclei (DAPI), microglia (Iba1), neurons (NeuN), astrocytes (GFAP), macrophages (CD68), and CD8^+^ T cells (CD8a) on left. Scale bar=100µm. Zoomed in image of nuclei (DAPI) and CD8^+^ T cells (CD8a) across groups in middle. Scale bar=20µm. Stacked bar plot showing quantification of the number of CD8^+^ T cells, CD4^+^ T cells, and B220^+^ B cells by age and VaD status on right. (D) UMAP embedding of T cells from murine SPLiT-seq showing *Gzmk* gene expression by age and treatment. (E) Representative flow cytometry gating strategy demonstrating surrogate markers used to identify splenic age-associated T cells (CD8a^+^ CD62L^−^ CD49d^+^ PD-1^+^ for 4-month-old and 22-month-old female C57BL/6N mice and quantification by age group (F). Mann-Whitney U-test. N=4/group. (G) Expression of *Gzmk* in CD8^+^ T cells from spleen of 4-month and 22-month-old female C57BL/6N mice by qPCR. N=4/group. Mann-Whitney U-test. Pseudotime trajectory analysis of murine brain SPLiT-seq CD8^+^ T_EFF_, CD8^+^ T_EM_, and CD8^+^ T_EM_ Gzmk clusters plotted onto StaVIA’s in-built VIA embedding demonstrating group (H), RNAvelocity showing the CD8^+^ T_EFF_ cluster as a less differentiated state compared to memory (T_EM_) CD8^+^ populations (I), and pseudotemporal projection of CD8^+^ T cells, showing that CD8^+^ T_EFF_ cells bifurcate into conventional CD8^+^ T_EM_ cells or CD8^+^ T_EM_ Gzmk cells (J). (K) Murine brain SPLiT-seq data showing *Gzmk*-expressing cells and the frequency that they co-express T cell or other immune cell markers with NK cell markers shown in pink. (L) Examination of *Gzmk* expression in aging spleen from publicly available bulk RNA-seq dataset (GSE132040). Statistical significance determined by one-way ANOVA with Tukey’s post-hoc test. N=4 mice/age group except 21-months which is N=3. (M) Ligand-receptor analysis of incoming signals to CD8^+^ T_EFF_, CD8^+^ T_EM_, and CD8^+^ T_EM_ Gzmk cells from our murine brain SPLiT-seq dataset showing ligand-receptor gene pairs, mean strength of interaction based on gene expression, and color depicting cellular source (ligand) sub-cluster. Multiple sources are shown for each receptor target on the top of the dotplot. N=3 biological replicates with 4-5 brains pooled per biological replicate for each group (total=52 brains). n=16,479 T cells.

We confirmed that the T cell cluster 1 (CD8 T_EM_ Gzmk) specifically expresses *Gzmk* compared to the other activated and memory CD8^+^ T cell clusters (CD8 T_EFF_ and CD8 T_EM_) and that *Gzmk* expression was specifically enriched at old age rather than young brains (Figure 5D). To confirm this, we performed flow cytometry using surrogate surface markers for the CD8 T_EM_ Gzmk cluster for live CD45^+^, CD3e^+^, CD8^+^, CD62L^−^, CD49d^+^, PD-1^+^ (Figure 5E). We confirmed that these cells were significantly enriched in aged brains compared to young brains (Figure 5F). We then sorted splenic CD8^+^ T cells from aged and young mice and confirmed that cells from the aged mice express markedly greater *Gzmk* expression by qPCR (Figure 5G). These findings indicate that aging is associated with the emergence of a distinct subset of antigen-experienced, pro-inflammatory CD8⁺ T cells that express *Gzmk*, suggesting a potential shift toward chronic cytotoxic and inflammatory T cell phenotypes in the aged brain.

To determine how the 3 differentiated CD8^+^ T cell clusters were related, we performed pseudotime and RNA velocity analysis on CD8^+^ T_EFF_, CD8^+^ T_EM_, and CD8^+^ T_EM_ Gzmk cells.^86^ Most of these cells were from the aged brains (Figure 5H). RNA velocity and pseudotime analysis reveal that the CD8 T_EFF_ cluster are less differentiated than either effector memory cluster (Figure 5I). CD8^+^ T_EFF_ cells either differentiate into the conventional CD8^+^ T_EM_ lineage or *Gzmk*-expressing CD8^+^ T_EM_ lineage (Figures 5I&J). Furthermore, the CD8^+^ T_EM_ Gzmk cluster is more terminally differentiated compared to the conventional CD8^+^ T_EM_ cluster (Figure 5I&J). This is in line with prior reports that CD8^+^ T_EM_ Gzmk are clonally expanded and terminally differentiated in disease-free aging in spleen, liver, lung, and peritoneum.^60,61^ This trajectory resembles the linear differentiation model, where naive CD8⁺ T cells differentiate through an early effector state before diverging into terminal effector or memory precursor fates.^87,88^ However, whereas Plumlee et al. and Abdullah et al. examined acute vesicular stomatitis virus infection over 1-2 week periods, our study examines CD8⁺ T cell differentiation in 3-month and 21-month naturally aged mice. Rather than identifying short-lived effector cells (SLECs) and memory precursor effector cells (MPECs), we observed long-lived differentiated CD8⁺ T cell subsets in the aged brain, with effector cells giving rise to both conventional CD8^+^ T_EM_ and *Gzmk*-expressing CD8^+^ T_EM_ memory lineages.

We next performed spectral flow cytometry of T cells from a cohort of female C57BL/6N mice. We examined the same four groups with n=5 mice in each control group and n=4 mice in each VaD group. Perfused, whole brains were harvested, and live, singlet T cells were identified as CD45^hi^ and either CD4^+^ or CD8^+^ (Figure S3A). T cell protein expression was examined by staining using a panel of antibodies targeting both naive and memory markers followed by PCA, Leiden clustering, and UMAP to identify T cell sub-populations (Figure S3B).^82,83^ We identified twelve distinct clusters of T cells which included naïve and memory CD4^+^ and CD8^+^ clusters of T cells as well as a population of B cells (Figure S3B&C).^44,82,83^ Similar to the scRNA-seq analysis, we found that the naïve T cell clusters (clusters 0 and 5) were more abundant in the young brains compared to aged groups (Figure 3M). We also found that the effector memory CD8^+^ T cells (CD8_EM and CD8_EM_CD28_PD1-int) were more abundant in aged brains compared to young groups (Figure S3D). We were unable to confirm by flow cytometry which cluster included the granzyme K-expressing CD8^+^ T cells because there are no commercially available monoclonal antibodies for murine granzyme K. However, we found that the CD8^+^ T_EM_ Gzmk scRNA-seq cluster of cells expressed *Cd28* and *Pdcd1* (encoding PD-1). We identified a similar cluster by spectral flow cytometry (cluster 3; CD8_EM_CD28_PD1-int) which had the same pattern as the granzyme-K-expressing cluster showing greater abundance in aged VaD brains compared to aged control brains (Figure S3D) similar to previous reports.^61^ These findings provide protein-level validation of our transcriptomic data, supporting the age-associated expansion of antigen-experienced memory CD8⁺ T cells in the brain in aged VaD mice.

There are reports that NK cells and other invariant T cell sub-types express granzyme K in humans.^56^ To examine this possibility in the murine brain, we examined *Gzmk*^+^ cells within our sc-RNAseq dataset. We identified whether T cell, NK cell, NKT cell, and invariant T cell genes were co-expressed with *Gzmk* (Figure 5K). We found that *Cd8a*, *Cd28*, *Il7r*, and *Cd44* had nearly 100% overlap with *Gzmk*-expressing cells (Figure 5K). Of the invariant T cell marker genes (identified in pink color), *Zbtb16* had the greatest overlap with *Gzmk*; however, an average of only 6% of *Gzmk*-expressing cells also expressed *Zbtb16* (Figure 5K). Furthermore, we wanted to determine at what age *Gzmk* expression begins to accumulate during the lifespan. As our dataset and other large sc-RNAseq datasets only include 3-months and 18-months but no ages between, we examined a publicly available bulk RNAseq dataset.^89^ We found that *Gzmk* expression begins to increase compared to 1- and 3-month old mice by 6-months and is significantly elevated by 12-months of age with a steady increase through 24-months of age (Figure 5L); however the increase from 9- to 12-, 15-, 18-, 21-, and 24-months of age was not statistically significant. This suggest that while *Gzmk* increases throughout the lifespan, the most appreciable increase is from 3-months to 12-months of age in mice (Figure 5L).

Finally, since most of the CD8^+^ T cells in aged brains are *Cd44*^+^ and thus appear to be antigen-experienced and activated or memory subsets, we wanted to explore the ligand-receptor interactions leading to their activation. To this end, we explored the ligand-to-receptor interactions between all sub-clusters to the three activated/memory T cell sub-clusters (CD8 T_EM_ Gzmk, CD8 T_EM_, and CD8 T_EFF_; Figure 5M). Most of the T cell inputs derived from microglial clusters with some inputs from myeloid cells (Figure 5M). Predominant ligands interacting with activated/memory T cell receptors were *Tgfbr1* ligands. There was also significant input from neutrophil *Hdc* to T cell *Hrh2*, *Hrh4*, and *Hrh1* receptors. Additionally, monocyte *Cd44* interacted with many different T cell receptors (Figure 5M). In general, most ligand receptor interactions were shared between all three activated/memory CD8^+^ T cell subsets with some notable exceptions. pDC *Apoe* to *Abca1* and neutrophil *Thbs1* to *Itga6* were unique inputs only to the CD8 T_EFF_ cluster (Figure 5M). Monocyte *Fn1* and *F13a1* to the CD8 T_EM_ Gzmk cluster *Itga4* were unique ligand-receptor interactions (Figure 5M). These unique interactions may be primary drivers of the fate decision of CD8 T_EFF_ cells to diverge either to conventional CD8 T_EM_ cells or the CD8 T_EM_ Gzmk subset.

### Distinct Microglial and T Cell Clusters Exhibit Specialized Secretory Roles in Cell-Cell Communication Networks

T cells respond to antigens presented and co-stimulation from antigen presenting cells (APCs).^90,91^ In the brain, microglia are the resident macrophages which can function as APCs under inflammatory conditions. Given the changes we observed in both microglial and T cell phenotypes with aging and VaD, we next asked whether there were transcriptional signatures of crosstalk between these two immune compartments. To examine potential signaling relationships between microglia and T cells in the brain in the context of aging and VaD, we performed network analysis of the microglia and T cell subclusters from our scRNA-seq dataset (Figure 6A). We identified the strength of all major outgoing and incoming signaling pathways between microglia and T cells based on receptor and ligand expression (Figures 6B&C). This revealed that CD8^+^ T_EFF_ cells participate heavily in both in sending and receiving signals, Homeostatic Microglia 1 participate mostly in receiving input, LDAMs participate heavily in outgoing signaling, and some clusters, such as Activated AP Microglia have limited participation in both (Figures 6B&C). Prior studies have demonstrated microglial can prime or restrain T cell activity depending on context.^92–94^

**Figure 6.**
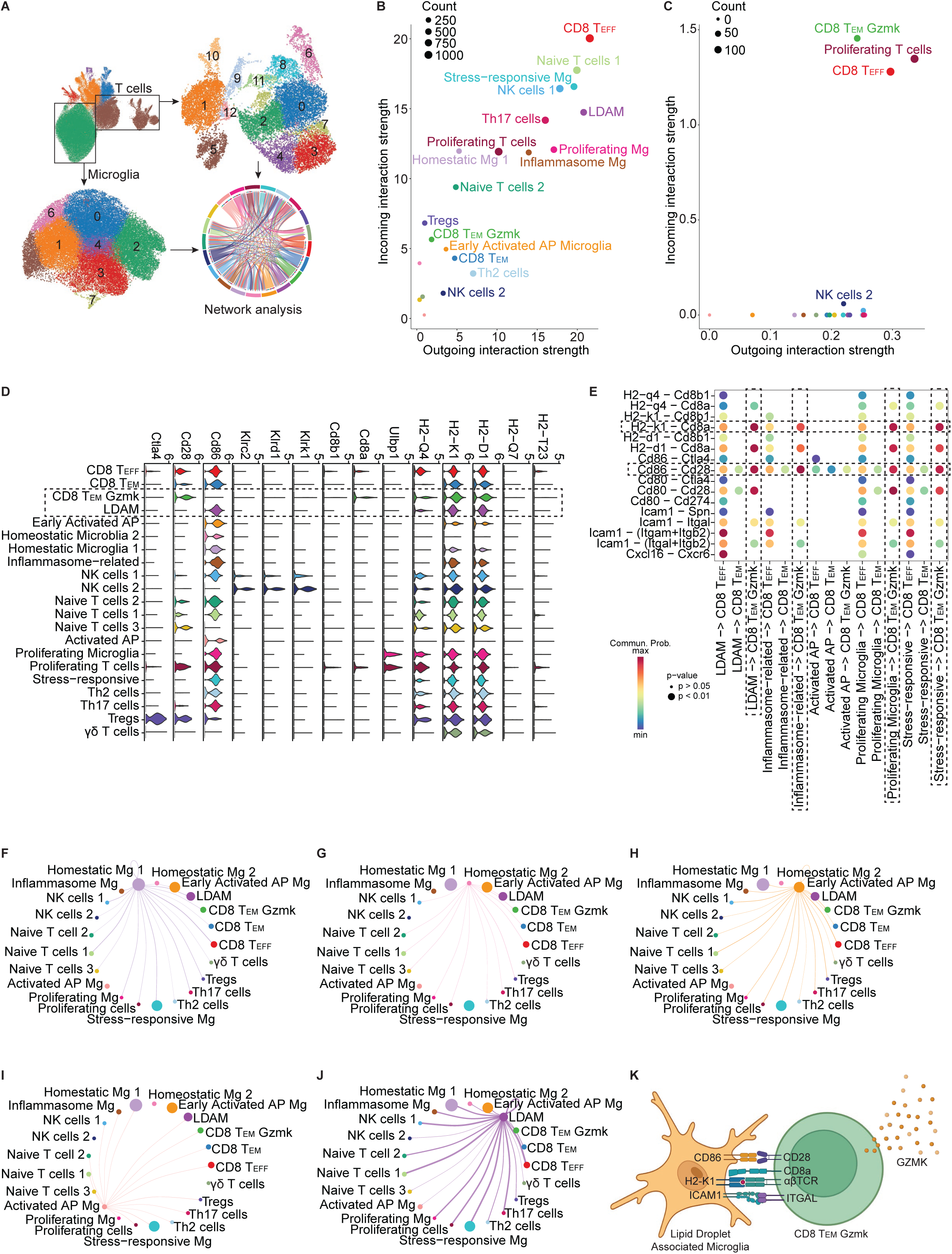
Microglia-T Cell Cross-Talk Drives Immune Dysregulation in the Aging Brain. (A) Workflow for cell–cell communication including Microglia and T cells using CellChat from the integrated single!Zcell RNA!Zseq dataset of young control, young VaD, aged control, and aged VaD mouse brains from our murine SPLiT-seq dataset. Hub analysis of all outgoing (x!Zaxis) versus incoming (y!Zaxis) interaction strength for all T cell and microglial clusters (B) and restricted to the CD86-CD28 signaling network (C). Bubble area denotes the number of significant ligand–receptor pairs. (D)LViolin plots showing normalized expression of representative ligand and receptor genes by cluster. (E)LDot plot of ligand-receptor pairs (y-axis) between microglial-to-t cell cluster pairs (x-axis). Rows = sending clusters; columns = receiving clusters. Dot diameter reflects significance and color reflects communication probability. Network plot of all outgoing signals to microglial and T cell sub-clusters from Homeostatic microglia 1 (F), Homeostatic microglia 2 (G), Early Activated AP microglia (H), Activated AP microglia (I), and LDAM (J). (K) Schematic of primary LDAM to CD8^+^ T_EM_ Gzmk cell-to-cell interactions. N=3 biological replicates with 4-5 brains pooled per biological replicate for each group (total=52 brains). n=36,900 microglial cells and n=16,479 T cells.

### *Gzmk*^+^ CD8^+^ T_EM_ Cells Accumulate with Age via CD86-CD28 Co-stimulation and MHC-I Signaling from Microglia

Prior literature suggests that microglial-to-lymphocyte co-stimulation and antigen presentation sustains neuroinflammation.^29,81,95^ Thus, we examined incoming and outgoing co-stimulatory and major histocompatibility complex class-I (MHC-I) signaling. We found complementary ligand-receptor signaling between the CD8 T_EM_ Gzmk cluster and LDAM clusters. We found the strongest signaling through the co-stimulatory molecule *Cd86*^96,97^ of several microglial clusters including the LDAM, inflammasome-related, proliferating, and stress-responsive microglial clusters with *Cd28* specifically on the CD8 T_EM_ Gzmk T cell cluster (Figure 6D&E). We also found the MHC-I^37^ genes *H2-K1* and *H2-D1* on the LDAM, inflammasome-related, proliferating, and stress-responsive microglial clusters corresponded to *Cd8a* on the CD8 T_EM_ Gzmk cluster (Figures 6D&E). We found that *Cd28* expression was present on all three activated/memory CD8 clusters but was most highly enriched in the CD8 T_EM_ Gzmk cluster. Within microglial clusters, the combination of *Cd86* and MHC-I genes were expressed among these four microglial clusters but were most enriched in the LDAM cluster (Figure 6D&E). These findings suggest a link between LDAMs either activate or sustain CD8 T_EM_ Gzmk cells in the aged brain. When comparing these microglial clusters’ outgoing signaling amongst other microglial clusters like Homeostatic microglia_1, we found that most of the microglial clusters signal widely to other microglial clusters and to most of the T cell clusters (Figures 6F-J). The LDAM cluster exhibited the strongest outgoing signaling network of all microglial clusters (Figure 6J). These results suggest that microglia, and LDAMs in particular, provide key signals to differentiate and sustain CD8 T_EM_ Gzmk cells within the aged brain environment (Figure 6K).

### Aging and VaD Enhance Myeloid Cell Accumulation in the Brain

We also identified significant shifts in myeloid cell clusters by age and treatment group with clusters 1 and 2 being more enriched in the aged brains (Figure S4A&B). Since the macrophage clusters were the most numerous myeloid cells, we examined significantly differentially expressed gene expression within the three macrophage clusters and performed pathway analysis (Figure S4C-E). We compared each macrophage cluster to the other macrophage clusters (e.g. cluster 0 compared with clusters 1 and 2 combined). In Macrophages_1 (cluster 0), which was enriched in young brains, we found differentially expressed genes related to interleukins 2, 4, 12, 13, 15 and 18 signaling (Figure S4C-E). In contrast, in Macrophages_2 (cluster 1) which was enriched in aged brains, especially in the aged VaD group, we found pathways related to metabolism, response to neutrophil degranulation, actin and cytoskeletal remodeling, and antigen presentation (Figure S4F-H). Finally, in Macrophages_3_PD-L1 (cluster 2), which was also enriched in aged brains compared to young, we found significantly enriched pathways related to cell adhesion, T cell activation, and interleukin 2 and 15 responses (Figure S4I-K). These findings suggest that macrophages with more interleukin secretion are predominant in young brains. At older ages, macrophages expressed transcriptional programs suggesting they have become metabolically more active and potentially activating local immune cells and T cells. Taken together, these results demonstrate that aging and VaD drastically remodel the myeloid compartment and likely contribute to the neuroinflammatory phenotype in the aging brain.

### Aging Influences Transcriptional Profiles of B cells in the Brain

B cells have previously been found to accumulate in the brains of aged mice.^76^ Thus, we also further examined the B cell sub-cluster in our scRNA-seq dataset (Figure S5A-B). When we group B cells by their treatment/age combination and analyze gene expression, regardless of cluster, we observe that expression of *Jchain* and high expression of BCR genes are enriched in the aged mice, regardless of treatment group (Figure S5C). Thus, our observations are consistent with previously published research indicating that ASCs can accumulate in the brains of aged mice. Given that both B1 cells and ABCs can rapidly differentiate into ASCs^77,98^, and ABCs accumulate in aged mice^76^, we asked the question whether we observed enrichment of this effector-like transcriptional profile in B cells in the brains of aged mice. When we examine B cells by age rather than cluster, we observe higher expression of ABC/B1/effector associated genes in the B cells of aged mice, including *Cd80*, *Zbtb20*, *Apoe*, *Ly6a*, and *Ahnak* (Figure S5C&D).^76,99^ Conversely, we observe enrichment of mature follicular B cell program genes like *Ebf1*, *Bach2*, *Fcer2a*, *Sell* and *Ighd* in the brains of young mice (Figure S5C&D).^72,100^ Consistent with these results we observe significant enrichment of clusters 0 and 2 in the brains of aged-control mice relative to young-control mice. We did not observe any differences between the treatment groups which indicates that VaD does not appear to affect accumulation of these cells in the brains of young or aged mice. Thus, the primary driver of differences in B cell populations was age rather than treatment, as we observed a small but significant increase in the frequency of ABC/B1-like B cells present in the brains of aged control mice, indicating that age may favor the accumulation of effector/innate-like B cells in the brains of aged mice. In total, these data support the notion that the brains of aged mice harbor transcriptionally distinct B cell populations compared to young mice.

## DISCUSSION

Here, we provide a comprehensive, validated immune cell compendium of the aging brain that complements prior aging^101^, immune aging^61^, and brain^26,37,38^ single-cell transcriptomic atlases. Previous studies using young animals did not capture T cells in the brain, our dataset integrates T cells, microglia, B cells, macrophages, and neutrophils from young and aged brains in both healthy and disease states. Here, we demonstrate significant shifts in immune cell populations, particularly microglia and T cells, in the context of age and VaD. In a recent paper which used intracranial injection of N5-(1-Iminoethyl)-L-ornithine dihydrochloride to induce subcortical ischemic lesions to model VaD, significant shifts in microglia from homeostatic toward pro-inflammatory phenotype similar to our findings.^8^ However, the animals in this study were young and T cells were not examined.^8^ Furthermore, our data are validated with subsequent spectral flow cytometry, where we identified shifts in both resident microglia and infiltrating peripheral CD8^+^ T cells. Among our key findings is the age-dependent accumulation of LDAMs, which are a specialized inflammatory subset of microglia characterized by immune-modulatory gene signatures. We also identified a distinct CD8^+^ T_EM_ Gzmk^+^ cells which had previously been identified in various compartments with aging but not yet in the brain.^56,57,61,62^ These are prominent hallmarks of immune aging which are enhanced in the context of VaD. This highlights their potential role in age-related neuroinflammation and neurodegeneration.

LDAMs exhibit extensive outgoing signaling pathways, strongly implicating them in the sustained inflammatory environment characteristic of VaD and other forms of neurodegenerative diseases.^46,48,102–104^ The age-dependent shift from homeostatic to pro-inflammatory LDAM underscores the critical role of age-related changes in microglia promoting neuroinflammation and cognitive decline. Our findings that both Early AP Microglia and Intermediate AP Microglia show a strong indication of declining homeostatic potential with increased pro-inflammatory and antigen-presenting potential. This shift toward an activated, antigen-presenting state likely perpetuates chronic inflammation and contributes to the age-dependent accumulation of effector and memory cytotoxic CD8^+^ T cell subsets in the brain. Our findings that CD8^+^ T_EM_ Gzmk^+^ cells express markers indicative of antigen experience, cytotoxicity, and exhaustion, such as *Cd44*, *Lag3*, *Tox*, *Pdcd1*, *Ccl5*, *Fasl*, *Eomes*, and *Tigit* supports this idea.^105,106^ Accumulation of CD8^+^ T_EM_ Gzmk^+^ cells correlates with enhanced co-stimulatory CD86-CD28 signaling and MHC-I-dependent interactions in the three activated microglial subsets that accumulated most with age.^17,32,91^ This is consistent with the emerging idea that chronic antigen exposure and inflammatory microenvironments in the aging brain promotes the differentiation and retention of cytotoxic, tissue-resident T cells.^24,25,107^

Despite their strong cytotoxic and inflammatory potential^57,108,109^, CD8^+^ T_EM_ Gzmk^+^ cells exhibited limited outgoing signaling capacity toward other T cell subsets or microglia compared to other differentiated CD8^+^ subsets, such as CD8^+^ T_EFF_ and traditional CD8^+^ T_EM_ cells. However, the strong incoming signals via the CD86-to-CD28 and MHC-I signaling from H2-d1 and H2-k1-to-CD8a, along with selective outgoing THY1 signaling, suggest a unique and possibly highly specialized functional role for CD8^+^ T_EM_ Gzmk^+^ cells. The presence of CD8^+^ T_EM_ Gzmk^+^ cells in aged VaD brains raises intriguing questions regarding their precise contributions to neuropathology especially given recent findings implicating this cell type in autoimmune diseases.^56–59,59,110–112^ In contrast the aging brain, these diseases all possess autoimmune or foreign body components (i.e., allogeneic transplant or infection). It is plausible that CD8^+^ T_EM_ Gzmk^+^ cells’ cytotoxic and inflammatory functions exacerbate neuronal injury and cognitive decline in age-related neurodegenerative diseases more broadly. A recent article also found CD8^+^ T_EM_ Gzmk^+^ cells in the brain in PS301^+/+^ tauopathy mouse model and human patients.^36^ This demonstrated CD8^+^ T_EM_ Gzmk^+^ cells in proximity to microglia and phospho-tau, and when PS301^+/+^ tauopathy mouse model was employed in *Cd8*^−/-^ mice, the loss of all CD8^+^ T cells enhanced phospho-tau deposition in the brain.^36^ The conclusion that granzyme K-expressing CD8^+^ T cells may be beneficial is similar to another recent report showing that CXCR6^+^ CD8^+^ T cells are brain-resident and perform immunoregulatory functions.^25^ Despite these two reports of more anti-inflammatory, immunoregulatory functions of granzyme K-expressing CD8^+^ T cells, a large body of work has demonstrated that CD8^+^ T_EM_ cells are largely pro-inflammatory in the brain^32–34,94,113–119^ and in many other disease contexts^56,57,59,109–111,120^ outside of infection and cancer. Taken together, our findings suggest that LDAMs, antigen-presenting microglia, and CD8^+^ T_EM_ cells all contribute to pro-inflammatory processes in aged brains in an interconnected fashion.^32,121^ Thus, therapeutic strategies aimed at restoring homeostatic microglial functions or resolving pro-inflammatory actions of LDAMs and activated microglial subsets could hold significant potential for management of neuroinflammatory diseases associated with aging.

Beyond the age- and VaD-dependent shifts in microglial and memory CD8^+^ T cell subsets and their relationships, our scRNA-seq dataset also reveals that age and VaD lead to shifts in other immune cell sub-types including CD4^+^ T cells, macrophages, and neutrophils. We observed the loss of naïve T cells and enrichment of activated CD4^+^ T cell subsets, and mature neutrophils in aged VaD brains. These cell types were not fully explored in this study but are clearly implicated in chronic adaptive immune activation and innate inflammatory processes driven by age and cerebrovascular pathology.^15,51,67^ Our scRNA-seq and spectral flow cytometry datasets serves as resources for the broader neuroscience and neuroimmunology communities to further examine the complex immune interactions in the aging brain. Leveraging these publicly accessible scRNA-seq and spectral flow cytometry datasets to explore the transcriptomic signatures and intercellular communication networks of all immune populations may be valuable resources to improve our understanding of neuroimmune contributions to age-related cognitive impairment and neurodegenerative diseases.

Limitations of our study include the lack of information on the origin of peripheral cells in the brain. The aged brains largely lacked naïve T and B cells but whether activated and memory T and B cells within the brain were derived first from naïve cells within the brain or differentiated prior to migration to the brain cannot be determined from our dataset. Future studies could employ lineage tracing models with aging to address these possibilities. Macrophages, neutrophils, T, and B cells are rare cell types in the brain relative to resident glia and neurons. Multiple brains were required to pool for each biological replicate to generate this atlas of brain-derived T cells. Thus, multiple batches of sorted cells from the brain were required for sufficient yield of single-cell RNA-seq which required fixation prior to library preparation and sequencing. A limitation of this approach is that TCR sequencing was not commercially available with SPLiT-Seq.^41^ The lack of data on T cell clonality is a limitation of our study, especially given prior findings that *Gzmk*-expressing CD8^+^ T_EM_ cells are the most clonally expanded compared with other T cell subsets.^19,60,61^ Since most of the lymphoid and myeloid cells are rare and multiple brains were pooled for each sample, we were unable to dissect specific brain regions in this dataset. As spatial transcriptomic methods increase in resolution with the ability profile the whole genome, future analyses could examine heterogeneity and localization at the same time.

Despite these limitations, we identified the emergence of LDAMs, CD8^+^ T_EM_ Gzmk^+^ cells, and dynamic neuroimmune interactions as defining features of immune aging in the brain. These findings significantly enhance our understanding of the immune mechanisms underlying vascular contributions to VCID and highlight microglia and CD8^+^ T cells as potential therapeutic targets for attenuating age-related neurodegenerative disease progression.

## EXPERIMENTAL MODELS AND SUBJECT DETAILS

### Data availability

All raw and processed, filtered, and annotated (.h5ad AnnData objects) scRNA-Seq data are available at the NCBI Gene Expression Omnibus GSE296215 with secure access token for reviewers (ofmdiekyrdqbzgj). Human microglia from healthy and VaD samples is publicly available at the NCBI Gene Expression Omnibus GSE213897.^84^ Murine aging spleen bulk RNAseq dataset is publicly available at the NCBI Gene Expression Omnibus GSE132040.^89^ Raw and processed, filtered, and annotated (.h5ad AnnData objects) spectral flow cytometry data are available at Figshare (Microglia: https://doi.org/10.6084/m9.figshare.30788516, and T cell: https://doi.org/10.6084/m9.figshare.30788582). Code used to analyze scRNA-Seq, spectral flow cytometry, and COMET imaging data are available upon request. Cell Analyzer for Flow Experiments (CAFE) was used to analyze spectral flow cytometry data (https://github.com/mhbsiam/cafe).^83^ Requests for additional data or materials can be made to the corresponding author at.

### Study approval

All animal experiments were carried out in accordance with the Institutional Animal Care and Use Committee at the University of Alabama at Birmingham (protocol no. 22627).

### Animal subjects

Female C57BL/6N wild-type (WT) mice aged 3 months (young) and 21 months (aged) were sourced from the National Institute on Aging (NIA) rodent colony and Charles River Laboratories (stock #027; Wilmington, MA). Mice were housed under specific pathogen-free conditions with a 12-hour light/dark cycle with free access of food and water. Cohort sizes were determined using power analyses (α = 0.05, β = 0.20) based on variance estimates from prior age-related immunity studies in this model.

### Bilateral Carotid Artery Stenosis Mouse Model of Vascular Dementia

Animals were anesthetized with inhaled isoflurane (2-3% in oxygen) and received a single subcutaneous injection of sustained-release buprenorphine (1 mg/kg) for analgesia. Titanium microcoils (Motion Dynamics Corp., part # MDC 29030A; wire diameter 0.18 mm inner diameter) were applied to both common carotid arteries (CCA). CCAs were exposed via midline cervical incision and gently isolated from surrounding tissues. Sterile 5-0 silk sutures were place under the CCA and used to lift the artery. Microcoils were placed around both CCAs just below the carotid bifurcation by holding the base of the microcoil and gently wrapping the artery around the coil using the suture.^43^ The suture filament was removed, vessel observed for normal flow, and the incision site was closed. Body temperature was maintained at 37°C throughout the procedure by operating on a heated surgical pad. Sham-operated mice underwent an identical procedure without microcoil placement. Following surgery (about 15 minutes from induction of anesthesia), mice recovered under close observation until fully ambulatory (typically about 20 minutes after cessation of anesthesia) and were monitored daily thereafter for three days. After surgery, investigators performing subsequent experiments were blinded to experimental groups.

### Isolation of Mouse Brain Cells

Brains were harvested from mice as previously demonstrated.^65,122^ Briefly, euthanasia and transcardial perfusion with 10-15 ml of ice-cold phosphate-buffered saline (PBS). The isolated brains were kept on ice in RPMI 1640 media supplemented with 10% fetal bovine serum (FBS), 1% penicillin/streptomycin, and 1% L-glutamine. The tissue was minced into small pieces and incubated at 37°C for 10 min with gentle rotation in 2 ml of enzyme mixture containing 1 mg/mL collagenase IV and 20 µg/mL DNase I. After initial incubation, the tissues were pipetted to break up large tissue fragments and incubated for an additional 10 min. Digested tissue was filtered through a 70 µm cell strainer (Fisherbrand) into a 50 mL conical tube and gently mashed using a 3 ml syringe plunger. Cells were washed with cold PBS (Ca^2+^/Mg^2+^-free, Gibco), and pelleted at 500 × g for 8 min at 4°C. The supernatant was discarded, and the pellet was resuspended in 7 mL of media and mixed with 3 ml of 90% Percoll, then gently vortexed, and transferred into a clean 15 mL tube. A 70% Percoll solution (2–3 mL) was gently layered below the cell suspension. Tubes were centrifuged at 500 × g for 32 min at 4°C with minimal acceleration and no braking. The cloudy buffy coat layer containing immune cells was carefully collected from the Percoll interface, washed with cold PBS, and centrifuged again at 500 × g for 8 min at 4°C. The pellet was resuspended in 500 µL cold PBS, filtered through a 70 µm cell strainer cap into a 5 mL flow cytometry (FACS) tube, and centrifuged at 500 × g for 5 min at 4°C and resuspended in FACS buffer (PBS/2% FBS/2 mM EDTA).

### Flow Cytometry and Cell Sorting of Murine Brain Immune Cells

Following cell isolation, single-cell suspensions were incubated with Live/Dead viability dye (1:100 dilution) in PBS for 5 min at 4°C, and subsequently blocked with Fc receptor blocking reagent (1:50 dilution in FACS buffer) for 10 min at 4°C. Cells were stained for 30 min at 4°C in the dark with the following fluorochrome-conjugated antibodies: anti-CD45 (BUV661, clone 30-F11, BD Biosciences, catalog no. 612975, 1 µL/sample), anti-CD3e (BB700, clone 145-2C11, BioLegend, catalog no. 152312, 1 µL/sample), anti-CD4 (BUV396, clone GK1.5, BioLegend, catalog no. 100491, 0.5 µL/sample), and anti-CD8a (APC-Cy7, clone 53-6.7, BioLegend, catalog no. 100766, 0.5 µL/sample). Stained cells were washed, centrifuged, and resuspended in 300 µL of FACS buffer. CD45^+^ CD3e^+^ cells were sorted using a FACSAria II cell sorter (BD Biosciences). Sorted cells were collected in cold FACS buffer and maintained at 4°C for fixation with Evercode Cell Fixation Kit v2.1.1 (Parse Bioscience).

### Library Preparation and Single-Cell RNA Sequencing of Murine Brain Immune Cells

Single-cell suspensions were fixed using the Evercode Cell Fixation Kit v2.1.1 (Parse Biosciences). Briefly, up to 150,000 cells per sample were centrifuged (200 ×g, 10 min, 4°C) in 15 mL polypropylene tubes using a swinging bucket rotor. Pellets were resuspended in 750 μL ice-cold Cell Prefixation Buffer supplemented with RNase Inhibitor, filtered through a 40 μm strainer, and transferred to fresh tubes. Fixation was initiated by adding 250 μL Cell Fixation Solution containing Cell Fixation Additive with immediate gentle mixing (3x pipetting), followed by 10 min ice incubation. Permeabilization was achieved using 80 μL Cell Permeabilization Solution (3 min ice incubation) before neutralization with 4 mL Cell Neutralization Buffer.

After centrifugation (200 ×g, 10 min, 4°C), pellets were resuspended in 150 μL ice-cold Cell Buffer containing RNase Inhibitor, filtered through a 40 μm strainer into 1.5 mL tubes, and maintained on ice. For cryopreservation, cells underwent stepwise DMSO addition (3 × 2.5 μL aliquots, 7.5 μL total) with gentle mixing between additions, followed by controlled freezing in a Mr. Frosty container (ThermoFisher Scientific) at -80°C. All resuspension steps used narrow-bore pipette tips to minimize doublet formation, with strict maintenance of ice-cold conditions throughout the protocol.

scRNA-seq libraries were prepared using the Parse Biosciences Evercode™ WT v2 kit (SKU: ECW02030) following the manufacturer’s protocol based on SPLiT-Seq technology.^41^ Briefly, fixed cells were subjected to combinatorial barcoding across four rounds to uniquely label transcriptomes. In Round 1, cells were distributed into a 48-well plate for reverse transcription using barcoded primers. Pooled cells then underwent two additional ligation-based barcoding rounds (Round 2: 96-well plate; Round 3: 96-well plate) to append unique adapters. Cells were lysed, partitioned into 7 sublibraries, and cDNA was amplified using template switching and PCR (12–14 cycles, depending on cell input). Amplified cDNA was fragmented, end-repaired, and A-tailed. Illumina adapters were ligated using the Parse UDI Plate - WT (UDI1001) for unique dual indexing (48 indices). Final libraries were size-selected using SPRI beads (KAPA Pure Beads) and sequenced on an Illumina NovaSeq 6000 with paired-end reads (Read 1: 66 cycles; i7/i5 indices: 8 cycles each; Read 2: 86 cycles).

### Cell Processing and Quality Control of Murine Brain Immune Cells

Cell viability and concentration were assessed via trypan blue staining and hemocytometer counting. Libraries were quantified using a Qubit™ Flex Fluorometer (ThermoFisher Scientific) and fragment size distribution validated on an Agilent 4200 TapeStation.

### scRNA-Seq Data Processing of Murine Brain Immune Cells

FASTQ files were processed using the Parse Biosciences computational pipeline (ParseBiosciences-Pipeline.1.3.1), aligning reads to the GRCm39 reference genome (Ensembl database), demultiplexing based on combinatorial barcodes, and generated gene expression matrix for each cell. To integrate and analyze scRNA-seq data derived from multiple samples, we employed the scVI (single-cell Variational Inference) model, a probabilistic deep learning framework designed to model the underlying structure of scRNA-seq data while correcting for batch effects and accounting for known sources of variation.^123^ Doublets were identified and removed using SOLO (scVI-tools v1.3.1), a deep learning-based method that probabilistically discriminates singlets from doublets by training a conditional variational autoencoder (cVAE) to model the likelihood ratio of observed vs. synthetic doublet gene expression.^124^ Cells with SOLO doublet scores exceeding an FDR-controlled threshold (q < 0.01, Benjamini-Hochberg correction) were excluded, achieving 100% recall of spiked synthetic doublets. Post-filtering, 69,614 high-confidence singlets were retained.

### Data Integration Single-cell RNA-seq Data Analysis with scVI of Murine Brain Immune Cells

Our dataset comprised 69,614 cells and 14,915 genes across 12 batches, corresponding to distinct 4 samples pooled per biological replicate with 3 biological replicates per group. All analyses were performed using scanpy (v1.9.5) and scvi-tools (v1.3.1) in Python (v3.11).^123,125^ Raw count matrices from 69,614 cells across 12 samples (3 biological replicates each of 22 month VaD, 22 month Control, 4 month VaD, and 4 month control), stored in adata.layers[’counts’], was used as the input for the scVI (Stochastic Cellular Variational Inference ) model. To prepare the data, we structured the AnnData object using scvi.model.SCVI.setup_anndata, designating the ‘sample’ column as the batch key to correct for sample-specific variations. Cell cycle variation was explicitly modeled by including expression levels of 85 canonical cell cycle genes (*Mcm5*, *Pcna*, *Cdk1*) as continuous covariates. This approach enabled the model to incorporate the expression of these genes as continuous variables influencing the generative process.

Single-cell RNA-seq data were analyzed using the scVI (single-cell variational inference) model implemented in the scvi-tools package (v1.1.6).^123^ The scVI model was configured with the following parameters: n_hidden=128, n_latent=10, n_layers=1, dropout_rate=0.1, dispersion set as ‘gene’, negative binomial (NB) gene likelihood, and a ‘normal’ latent distribution. We adopted the NB count likelihood rather than the zero-inflated negative binomial (ZINB), as recent evidence indicates that NB sufficiently captures scRNA-seq data variability with reduced model complexity.^126^ Batch effects arising from different experimental samples were corrected by including ‘sample’ as a batch covariate, while cell cycle-related variability was explicitly modeled using 85 established cell-cycle genes as continuous covariates. Model training was conducted for 115 epochs. Latent embeddings obtained from the trained model were stored as X_scVI in the AnnData object and subsequently utilized for dimensionality reduction, clustering, and visualization. Normalized expression values scaled to a library size of 10,000 counts per cell were computed and stored that enabled cell-type specific differential expression analyses.

### Dimensionality Reduction and Clustering of Murine Brain Immune Cells

We performed data normalization and dimensionality reduction using the scVI framework (v1.0.0) to learn a latent representation of the single-cell transcriptomic data. This latent space captures the biological variability of the data while mitigating technical artifacts such as batch effects. Normalized gene expression values were computed from the trained scVI model. For clustering, we constructed a neighborhood graph using the scanpy.pp.neighbors function from the Scanpy toolkit.^125^ The latent space (X_scVI, 10 dimensions) was used as input for downstream analyses. A neighborhood graph was computed using 50 nearest neighbors in the latent space, followed by UMAP embedding (min_dist=0.5) and Leiden clustering (resolution=0.5) for unsupervised cell grouping and we got 16 unique clusters.^44,127,128^ Although principal component analysis (PCA) was performed on highly variable genes using scanpy.tl.pca with 50 components, this step was not utilized in the subsequent clustering or visualization workflows, which relied exclusively on the scVI latent representation.

### Differential Gene Expression Analysis of Murine Brain Immune Cells

Differential gene expression across identified clusters was evaluated using two complementary methods. First, Scanpy’s Wilcoxon rank-sum test was employed via sc.tl.rank_genes_groups to identify cluster-specific markers (parameters: groupby=’leiden’, method=’wilcoxon’, n_genes=20, use_raw=False). Second, the trained scVI model’s built-in Bayesian differential expression method was applied using model.differential_expression(groupby=’leiden’).^129^ We also performed pseudo-bulk differential expression pipeline using decoupler (v.1.8.0) and pydeseq2 (v.0.4.12). After aggregating raw counts for each cluster–sample combination using decoupler.get_pseudobulk, we filtered lowly expressed genes and built a pydeseq2 (DESeq2-like) model for each cluster comparison of interest. We generated effect size estimates as log_2_ fold changes and statistical significance which we plotted with volcano plots. We then performed over-representation analysis with MSigDB gene sets (including gene ontology biological processes and reactome collections) using decoupler functions (get_ora_df) to identify significantly enriched pathways for each cluster comparison. Results from all methods were integrated to characterize transcriptional patterns defining distinct immune cell populations in the aging brain.

### Cell Type Annotation of Murine Brain Immune Cells

For cell type annotation, murine single-cell RNA-seq data were clustered using the Leiden algorithm, with expression values scaled and thresholded to optimize population resolution. We used the canonical markers for identifying the cell types. We identified T cells (*Cd3e*, *Itk*, *Themis*), Microglia (*Tmem119*, *P2ry12*, *Lpl*, *Apoe*)^49^, Macrophages (*Mertk*, *Cd74*, *Lyz2*, *Csf1r*), Neutrophils (*S100a8*, *S100a9*, *Hdc*), and neurons (*Nkain2*, *Ptprd*, *Slc24a2*). We further sub-clustered microglia, T cells, and macrophages/neutrophils then performed Leiden clustering again, followed by cell type annotation by additional marker genes.

We further subset the T cells based on the canonical marker (*Cd3e*). We performed subset analysis on 16,479 T cells isolated from original dataset to resolve T cell heterogeneity. Raw counts (14,915 genes) were filtered to retain genes detected in ≥10 cells (12,158 genes remaining), ensuring robust feature selection. We used scvi-tools (v1.1.6.)^123^, we adapted our established scVI integration workflow previously applied to all immune cells with T cell specific optimizations. The model was trained on the ‘counts’ layer using ‘sample’ as batch covariate while controlling for technical variation (n_genes_by_counts, total_counts, pct_counts_mt). A two-layer neural network architecture with negative binomial likelihood was implemented and trained for 400 epochs to stabilize latent representations. From the trained model, we derived a 10-dimensional latent space (adata.obsm[’X_scVI’]) and computed a k-nearest neighbor graph (k=50) for Leiden clustering at resolution 0.5, yielding 12 transcriptionally distinct clusters which were identified using information from us and others.^60,61^ CD8^+^ T cell subsets were defined as follows: cluster 0 (CD8^+^ T_EFF_) by *Gzmb* and *Prf1*; cluster 1 (CD8^+^ T_EM_ Gzmk) by elevated *Gzmk* and *Tox*; cluster 2 (CD8^+^ T_EM_) by *Ccr5* and low *Sell*. Naive T cells (clusters 3–5) expressed *Sell* (Cd62L), *Ccr7*, and *Tcf7*. Th2 (cluster 6) showed *Gata3* and *Il4*; Th17 (cluster 8) expressed *Rora* and *Il23r*. γδT cells (cluster 9) were identified by *Trdc*, *Themis*, and lack of *Cd8a* and *Cd4*. Tregs (cluster 12) expressed *Foxp3* and *Il2ra*. NK cells (clusters 7, 10) were defined by *Ncam1* (CD56), *Klrb1c*, *Klrd1*, and *Gnly*. Proliferating cells (cluster 11) exhibited *Mki67* and *Top2a*.

These identifications were found using differential gene expression analysis for marker genes of specific T cell sub-clusters compared to all other T cells. After initial clustering, the microglial metacluster was further subset (n = 39,600 cells) and reanalyzed with the same scVI tools framework. We identified homeostatic microglia (clusters 0 and 6) by *Sall1*, *Fcrls*, and *P2ry13*, while Lipid-droplet Associated Microglia (LDAM, cluster 3) expressed *Apoe* and *Clec7a*. Stress-responsive microglia (cluster 2) expressed *Fos* and *Hspa1a*, and Inflammasome-related microglia (cluster 4) expressed *Casp4* and *Nlrp3*.

Proliferating microglia (cluster 7) expressed *Bub1b*. Early activated AP microglia (cluster 1) expressed *Ezh2,* and Activated AP microglia (cluster 5) expressed *Nfkb1* and *Lyve1*. These identifications were found using differential gene expression analysis for marker genes of specific microglial sub-clusters compared to all other microglia.

After initial clustering, the myeloid metacluster was further subset (n = 7,913 cells) and reanalyzed with the same scVI tools framework. We identified macrophages (clusters 0, 1 and 2) by *Mertk*, *Adgre1*, and *Cd86*. We identified pDCs (cluster 3) by lack of *Adgre1*, *Mertk*, *Itga4*, and lower *Itgax* and expression of *Cd83*, *Sirpa*, and *Plxdc2*. We identified Monocytes (cluster 4) by lack of *Mertk* and *Adgre1* and expression of *Itga4*, *Itgb2*, and *Cd86*. We identified Proliferating cells (cluster 5) by high expression of DNA replication and chromatin assembly genes. We identified cDC1s (cluster 6) and cDC2s (cluster 7) by low expression of *Mertk* and *Adgre1* and expression of *Itga4*, *Il31ra*, *Cd86*, *Cd83*, *Mef2c*, and *Il1b*. We differentiated these 2 based on expression of *Cd74* and MHC-II gene expression in cDC2s. Finally, we identified Neutrophils (cluster 8) by lack of expression of macrophage and monocyte markers and positive expression of *S100a8* and *S100a9*. These identifications were found using differential gene expression analysis for marker genes of specific microglial sub-clusters compared to all other myeloid cells, microglia, and T cells.

### Cell-Cell Communications Analysis of Murine Brain Immune Cells

Intercellular communication networks were analyzed from scRNA-seq data using CellChat (v1.6.0)^85^ and the complete CellChatDB, excluding non-protein signaling interactions to focus on secreted immune mediators. Communication probabilities were computed via computeCommunProb using the triMean method to calculate average ligand/receptor expression per cell group. Signaling dynamics were systematically mapped by integrating network theory, identifying dominant ligand-receptor interactions and functionally coordinated cell-cell communication events through integrative network analysis and pattern recognition.

CellChat objects were created from expression data (createCellChat). Overexpressed genes (p < 0.05, hypergeometric test) and ligand–receptor pairs (FDR < 5%) were identified (identifyOverExpressedGenes, identifyOverExpressedInteractions). Communication probabilities were calculated (computeCommunProb, filterCommunication, computeCommunProbPathway), and significant interactions were identified by comparing observed probabilities to those from 1,000 permutations of randomized cell labels. Interaction networks were then summarized (aggregateNet), and centrality metrics were computed (netAnalysis_computeCentrality). Dominant signaling pathways and ligand–receptor pairs were visualized using netVisual_bubble and netVisual_circle. Pathway-specific communication patterns (e.g., CXCL, TNF) were visualized using netVisual_circle and netVisual_chord, while global interaction weights and counts were mapped via netVisual_heatmap. Outgoing/incoming signaling modules were identified using non-negative matrix factorization (NMF) that will allow us to determine the number of patterns in outgoing and incoming signal based on each cell type. These patterns were validated through river (netAnalysis_river) and dot plots (netAnalysis_dot). All analyses were parallelized (30 workers) to ensure computational reproducibility. Major signaling inputs/outputs and functional coordination between cell populations were resolved through this framework.

### Pseudotime Trajectory Analysis and RNAvelocity of Murine Brain Immune Cells

Single-cell RNA velocity and pseudotime trajectories for microglia were analyzed using scVelo (v0.3.3).^79^ First, moments of spliced/unspliced counts were computed using PCA (10 principal components) and 10 nearest neighbors. Stochastic RNA velocities were then estimated to model transcriptional dynamics. A velocity-embedding graph was constructed to project trajectories onto the UMAP embedding. Velocity pseudotime was calculated by these velocities through the trajectory graph. Streamplots and pseudotime values were visualized using Scanpy’s UMAP embedding with Leiden cluster annotations and a gnuplot colormap, respectively.

Single-cell RNA velocity and pseudotime trajectories for CD8^+^ T effector and memory T cell clusters were performed on T cell clusters 0, 1, and 2 using pyVIA (v2.0.0).^86^ Trajectory inference was initiated from a root cell (index=2697). Terminal states were identified based on betweenness centrality and out-degree. Pseudotime and lineage likelihoods were projected onto a 2D embedding computed using VIA’s built-in StaVia embedding with min_dist=0.4. Trajectory curves were visualized on the resulting embedding.

### Spectral Flow Cytometry Data Analysis of Murine Brain Immune Cells

Single-cell suspensions were stained with two optimized spectral flow cytometry panels for microglia and T cell characterization, using a BD FACSymphony A5 SE Spectral Flow Cytometer (BD Biosciences) with BD FACSDiva software version 9.0. Initial data processing was done with FlowJo version 10.10. Cells were pre-treated with TruStain FcX™ PLUS (anti-mouse CD16/32, clone S17011E; BioLegend, Cat# 156604, Lot# B362118) to block Fc receptors. All antibodies were titrated for optimal signal-to-noise ratios. Data were analyzed with CAFE (v2.0.2) following our previously described methods.^82,83^ Antibodies used for flow cytometry staining:

T Cell Panel:

Live/Dead Blue dye (ThermoFisher, Cat# L23105)

CD4 Spark UV387 (clone GK1.5; BioLegend, Cat# 100492, Lot# B388730)

CD27 BUV615 (clone 37.51; BD Biosciences, Cat# 751529, Lot# 3291617)

CD45 BUV661 (clone 30-F11; BD Biosciences, Cat# 612975, Lot# 2174908)

TCRγδ BUV736 (clone GL3; BD Biosciences, Cat# 748997, Lot# 3303650)

PD-1 BV421 (clone 29F.1A12; BioLegend, Cat# 135213, Lot# B367777)

LAG3 Superbright 436 (clone eBioC9B7W; Invitrogen, Cat# SB436-1720)

KLRG1 BV510 (clone 2F1/KLRG1; BioLegend, Cat# 138421, Lot# B395543)

CD62L BV605 (clone MEL-14; BioLegend, Cat# 104438, Lot# B386347)

MHC II (I-A/I-E) BV711 (clone M5/114.15.2; BioLegend, Cat# 107643, Lot# B377900)

CD45R/B220 BV786 (clone RA3-6B2; BD Biosciences, Cat# 563894, Lot# 3150973)

CD49d FITC (clone R1-2; Invitrogen, Cat# 11-0492-82, Lot# 2796430)

CXCR3 PE-eFluor610 (clone CXCR3-173; Invitrogen, Cat# 61-1831-82, Lot# 2518430)

CCR5 PerCP-eFluor710 (clone HM-CCR5(7A4); Invitrogen, Cat# 46-1951-82, Lot# 2487957)

MHCI (H-2Kb/H-2Db) PE-Cy7 (clone 28-8-6; BioLegend, Cat# 114616, Lot# B406395)

CXCR6 APC (clone SA051D8; BioLegend, Cat# 151207, Lot# B363617)

CD44 Alexa Fluor700 (clone IM7; BioLegend, Cat# 103026, Lot# B370174)

CD8a APC-Fire750 (clone 53-6.7; BioLegend, Cat# 100766, Lot# B363617)

CD28 BUV805 (clone 37.51; BD Biosciences, Cat# 752545, Lot# 3291629)

CD3ε BV570 (clone 145-2C11; BD Biosciences, Cat# 563024, Lot# 4127753)

Microglia Panel:

Live/Dead Blue dye (ThermoFisher, Cat# L23105)

Ly-6C UV387 (clone HK1.4; BioLegend, Cat# 128060, Lot# B410594)

CD45 BUV661 (clone 30-F11; BD Biosciences, Cat# 612975, Lot# 2174908)

Siglec H BUV805 (clone 551; BD Biosciences, Cat# 752584, Lot# 4131860)

CCR2 BV421 (clone 475301; BD Biosciences, Cat# 747963, Lot# 4131859)

LAG3 Superbright 436 (clone eBioC9B7W; Invitrogen, Cat# SB436-1720)

CD11b BV510 (clone m1/70; BD Biosciences, Cat# 612800, Lot# 4127753)

CD3ε BV570 (clone 145-2C11; BD Biosciences, Cat# 563024, Lot# 4127753)

CD64 BV605 (clone X54-5/7.1; BioLegend, Cat# 139323, Lot# B412183)

MHC II (I-A/I-E) BV711 (clone M5/114.15.2; BioLegend, Cat# 107643, Lot# B377900)

CD11c BV786 (clone N418; BioLegend, Cat# 117335, Lot# B401468)

Ly-6G FITC (clone 1A8; BioLegend, Cat# 127606, Lot# B400511)

CXCR3 PE-eFluor610 (clone CXCR3-173; Invitrogen, Cat# 61-1831-82, Lot# 2518430)

CCR5 PerCP-eFluor710 (clone HM-CCR5(7A4); Invitrogen, Cat# 46-1951-82, Lot# 2487957)

MHCI (H-2Kb/H-2Db) PE-Cy7 (clone 28-8-6; BioLegend, Cat# 114616, Lot# B406395)

CD63 APC (clone NVG-2; BioLegend, Cat# 149006, Lot# B365325)

CX3CR1 Alexa Fluor700 (clone SA011F11; BioLegend, Cat# 149036, Lot# B409648)

CD38 APC-Fire750 (clone 90; BioLegend, Cat# 102738, Lot# B418109)

Trem2 APC (clone 6E9; BioLegend, Cat# 824806, Lot# B426400)

### High-Plex Spatial Phenotyping via Sequential Immunofluorescence of Murine Brain

High-plex spatial phenotyping was performed using the COMET™ automated platform (Lunaphore Technologies, version 1.1.0.0). We utilized a Sequential Immunofluorescence (SeqIF) approach to generate a multi-dimensional protein atlas from single 5µm coronal brain sections from the mouse brain. This method overcomes the spectral limitations of conventional microscopy by iteratively staining, imaging, and eluting antibodies from the same tissue section to generate a multi-dimensional protein atlas from a single sample. We analyzed coronal brain sections from four experimental cohorts processed in parallel: aged and young C57BL/6N mice subjected to BCAS or sham surgery.

The automated protocol executed 21 discrete cycles over approximately 40-42 hours. Each cycle followed temperature-controlled (37°C) regimen: primary antibody incubation (8 min), detection with fluorophore-conjugated secondary antibodies (4 min), DAPI nuclear counterstaining (ThermoFisher 62248, 1:1000, 2 min), multi-channel image acquisition across a 12.5 × 12.5 mm field-of-view, antibody elution (Elution Buffer BU07-L, Lunaphore, 2 min), and quenching (Quenching Buffer BU08-L, Lunaphore, 30 s). All washing steps utilized Multistaining Buffer (BU06, Lunaphore). The sequencing strategy incorporated quality controls throughout, including autofluorescence baseline acquisition, secondary-only negative controls at multiple intervals, and lectin-based controls at the protocol terminus to monitor elution efficiency and non-specific binding.

The 19-marker immunophenotyping antibodies and reagents are listed in Supplemental Table 1 and included: CD11c (Cell Signaling Technology 97585, D1V9Y, rabbit, 1:200), Foxp3 (Invitrogen 14-5573-82, FJK-16s, rat, 1:100), CD206 (Cell Signaling Technology 24595, E6T5J, rabbit, 1:200), CD4 (Invitrogen 14-9766-82, 4SM95, rat, 1:200), CD8 (Invitrogen 14-0195-82, 4SM16, rat, 1:200), NeuN (Abcam ab177487, EPR12763, rabbit, 1:2000), CD31 (Cell Signaling Technology 77699, D8V9E, rabbit, 1:200), CD3 (Abcam ab11089, CD3-12, rat, 1:200), myeloperoxidase (two distinct clones: Invitrogen MA1-80878, 2C7, rabbit, 1:100 and Abcam ab9535, polyclonal, rabbit, 1:50), Granzyme K (Invitrogen PA5-50980, polyclonal, rabbit, 1:200), GFAP (Abcam ab68428, EPR1034Y, rabbit, 1:200), APOE (Abcam ab183597, EPR19392, rabbit, 1:2000), CD45 (Abcam ab10558, polyclonal, rabbit, 1:200), Iba-1 (Abcam ab178846, EPR16588, rabbit, 1:1000), B220/CD45R (Invitrogen 14-0452-85, RA3-6B2, rat, 1:400), Arginase-1 (Invitrogen PA5-29645, polyclonal, rabbit, 1:500), CD44 (Cell Signaling Technology 37259S, E7K2Y, rabbit, 1:200), CD74 (Cell Signaling Technology 82174S, F3R3L, rabbit, 1:400), CD11b (Abcam ab133357, EPR1344, rabbit, 1:5000), Na+/K+ ATPase α (Santa Cruz SC-48345, H-3, mouse, RTU), CD68 (Abcam ab283654, EPR23917-164, rabbit, 1:100), and CoxIV (Abcam ab16056, polyclonal, rabbit, 1:200). The protocol concluded with 2 additional lectin stains: Maackia Amurensis Lectin I (Vector Laboratories FL-1311-2, 1:100) and Sambucus Nigra Lectin (Vector Laboratories CL-1305-1, 1:1000).

Secondary antibodies included goat anti-rabbit CoraLite Plus 555 (Proteintech RGAR003, RTU), goat anti-rat Alexa Fluor Plus 647 (Life Technologies A48265, 1:200), goat anti-rabbit CoraLite Plus 647 (Proteintech RGAR005, RTU), and goat anti-mouse CoraLite Plus 555 (Proteintech RGAM003, RTU). Antibody pairs were strategically combined to maximize signal separation and minimize cross-reactivity. Imaging was performed with optimized exposure times across all channels (DAPI: 25 ms; TRITC: 250 ms; Cy5: 400 ms; FITC: 500 ms; Cy7: 1000 ms) to enable quantitative comparison.

Following image acquisition, raw data were processed using HORIZON^TM^ viewer software (v-2.3.0.0) and Visiopharm software (v-2025.08.1.18881 x64) and exported as single-channel images for downstream analysis in ImageJ. Microglial regions of interest (ROIs) were defined using the Iba-1 channel thresholding. To generate cell masks, Iba-1 images were binarized, followed by morphological operations including two iterations of dilation and closing to consolidate cellular boundaries. The ‘Analyze Particles’ function was employed to identify cells, filtering for objects to include pixel size of >500 and <100,000 to exclude debris and non-specific noise. The resulting microglial ROIs were saved to the ROI Manager and applied across all spatially registered phenotypic marker channels (e.g., APOE, CD68, CD74, Arginase-1). Mean Fluorescence Intensity (MFI) was measured for each marker within each defined single-cell ROI. Data were exported as CSV files containing single-cell barcodes, sample metadata, and expression values for the complete antibody panel.

Single-cell MFI data were imported into Python and analyzed using the Scanpy toolkit. Data were log-normalized (scanpy.pp.log1p) to stabilize variance. PCA was used to reduce dimensionality, followed by the construction of a neighborhood graph (scanpy.pp.neighbors). Two-dimensional embeddings were generated using UMAP to visualize cellular heterogeneity. Unsupervised clustering was performed using the Leiden algorithm (resolution = 0.4) to identify distinct microglial states. Cluster identities were annotated based on the expression of canonical and state-specific markers (e.g., CD206, APOE, Arginase-1, CD11b, CoxIV) visualized via dot-heatmap plots and UMAP overlays. The frequency of cells per Leiden cluster was quantified and compared across experimental groups (Aged/Young, BCAS/Sham).

To quantify spatial interactions, cell centroid coordinates and identities were extracted from fluorescence images using ImageJ. ImageJ-derived coordinate tables were imported into Python and analyzed using pandas and numpy libraries, with nearest-neighbor calculations performed using the cKDTree algorithm from scipy.spatial. For each image, unique cell types were enumerated, and pairwise nearest-neighbor distances were computed by constructing KD-trees for each target population and querying the closest target cell to every source cell (k = 1).

Resulting distances, source and target IDs, and cell-type information were consolidated across four group-specific datasets for quantitative comparison. Resulting distances, source and target IDs, and cell-type information were consolidated across four group-specific datasets for quantitative comparison.

### Human Single-Nucleus RNA-Seq Data Acquisition and Analysis

Publicly available single-nucleus RNA-sequencing (snRNA-seq) data from human periventricular white matter were obtained from the Gene Expression Omnibus (GEO; GSE213897), as originally described.^84^ This dataset comprises samples from patients with VaD and age-matched normal controls (NC). The ERG-sorted (DAPI+ERG+) nuclear fraction, enriched for microglial populations, was utilized for the present analysis. Raw count matrices, barcodes, and feature files for each sample were parsed into AnnData objects. Mitochondrial, ribosomal, and hemoglobin genes were annotated based on gene symbols to enable computation of standard QC metrics (gene counts, UMI counts, and mitochondrial transcript percentages). Cells with <150 genes or >5% mitochondrial transcripts were filtered out, and genes expressed in <3 nuclei were excluded.

Raw counts were normalized (sc.pp.normalize_total and sc.pp.log1p). Highly variable genes (HVGs) were identified with scanpy.pp.highly_variable_genes using the Seurat v3 flavor, 4000 features while accounting for batch structure via the per!Zsample GSM identifier. Counts were scaled (sc.pp.scale), and PCA was performed (50 components). Neighborhood graphs were constructed using 30 principal components and 15 nearest neighbors per cell (sc.pp.neighbors), followed by UMAP for two!Zdimensional visualization. Unsupervised clustering was performed by Leiden (0.5 resolution) to define transcriptionally distinct microglial clusters

Batch effects were identified by insufficient group cell type integration into Leiden clusters across samples. Thus, Harmony integration was applied to the PCA embedding using a composite batch covariate combining GSM identifier and marker status (ERG vs DAPI; sce.pp.harmony_integrate) which converged after 3 iterations. The Harmony!Zcorrected principal components were used to recompute neighborhood graphs, UMAP embeddings, and Leiden clusters, yielding integrated microglial populations in which samples and markers were well mixed across clusters.

The Leiden clusters containing microglial nuclei were identified by expression of *CX3CR1*, *ITGAM*, *TMEM119*, *APOE*, *TREM2*, and absence of *OLIG2* or *GFAP*. No T cells or neurons were identified in this dataset. The microglia were analyzed by re-computing the nearest neighbors, UMAP, and Leiden (resolution=0.8). Cluster identities were assigned based on differential expression of established microglial signature genes, including homeostatic markers (*TMEM119*, *P2RY13*, *MAFB*, *SALL1*), activation markers (*APOE*, *CLEC7A*, *LPL*, *ITGAX*), inflammatory mediators (*IL1B*, *IL1A*, *TNF*, *CCL3*, *CCL4*), and stress-response genes (*HSP90AA1*, *DNAJB1*, *EGR1*). UMAP embedding was performed for visualization.

To identify conserved microglial transcriptional responses between human VaD and murine VaD, differentially expressed genes were identified for both species. Mouse genes were mapped to human orthologs, and the overlap between species-specific VaD gene signatures was visualized using Venn diagrams. Pseudo-bulk expression profiles were generated for selected microglial marker genes across conditions for both species. Expression values were z-score normalized and subjected to K-means hierarchical clustering. Cluster proportions were compared across mouse conditions (4-month control, 4-month VaD, 22-month control, 22-month VaD) and human conditions (healthy control and VaD).

### Bulk Murine Splenic Aging Analysis of Granzyme K Expression

Murine splenic aging bulk RNAseq data (GSE132040) were exported directly from the web-hosted shiny app (https://twc-stanford.shinyapps.io/maca/)^89^ and imported directly into GraphPad Prism for figure generation and data analysis.

### Statistical Methods

Quantification and statistical analyses were performed using GraphPad Prism (v10.4.1). All results are presented as meanL±Ls.e.m. Normality was determined using the Shapiro–Wilk test. Non-parametric tests were used for data that are not normally distributed or when groups contained fewer than eight biological replicates. Data with one independent variable (that is, age) were analyzed using two-tailed Student’s t-test or two-tailed Mann–Whitney U-test (non-parametric). When more than one independent variable was analyzed (that is, age and treatment or repeated measures), one-way ANOVA (or Kruskal–Wallis test) or two-way ANOVA with multiple comparisons were used. Differential gene expression analysis of single-cell RNA sequencing data was conducted using Wilcoxon rank-sum test, along with a Bayesian inference method provided by a trained scVI model. Two-sided P values were used, and P values less than 0.05 were considered significant except where correction for multiple comparisons was performed (such as single-cell RNA-seq analysis). Details regarding specific analytical tests used are provided in the figure legends.

## CONTRIBUTIONS

M.A.A. and D.J.T. conceived the project. M.A.A., M.H.B.S., D.V.III, C.B., J.N.B., A.W., C.R.R., H.T., J.S., A.N.H., M.P., and D.J.T. analyzed the data. M.A.A. and D.J.T. wrote the paper. All authors reviewed and edited the paper.

## ETHICS DECLARATIONS

The authors have no competing financial conflicts of interest.

## ACKNOWLEGMENTS

This study was supported by National Institutes of Health (NIH) awards AG068309 (D.J.T.), the UAB Nathan Shock Center, which is supported by the National Institutes of Health/National Institute on Aging Grant P30 AG050886. This research was conducted while D.J.T. was an AFAR Grant for Junior Faculty awardee. The funders had no role in study design, data collection and analysis, decision to publish, or preparation of the manuscript. We thank the staff at the UAB Flow Cytometry and Single Cell Core laboratory which is supported by the Center for AIDS Research, AI027767 and the O’Neal Comprehensive Cancer Center, CA013148. The BD FACSymphony A5 SE spectral flow cytometer at UAB was purchased under S10 OD032296 (to Troy Randall). We acknowledge the Heflin Center for Genomic Sciences at UAB for sequencing data. We acknowledge the UAB high-performance computing support and CPU time on the Cheaha compute cluster.

---

**Figure S1. Gating Strategy for Sorting Immune Cells from the Brain**

Flow-assisted cell sorting (FACS) of immune cells from the murine brain to isolate cells, singlets, live cells, CD45^int^ and CD45^hi^ resident immune cells, peripheral myeloid cells and CD3^+^ T cells followed by confirmation of CD8 and CD4 expression within the CD3^+^ T cells prior to performing barcoding for SPLiT-seq analysis.

**Figure S2. Spectral Flow Cytometry Corroborates Murine Brain Microglial Heterogeneity in Aging and Vascular Dementia**

(A) Gating strategy for identifying live singlet microglia as CD45^+^ CD11b^+^ CX3CR1^+^ CD38^−^ cells. This population was then analyzed using principal components analysis (PCA), uniform manifold approximation projection (UMAP), and Leiden clustering. (B) Microglia cells shown by UMAP embedding split by age and treatment depicting Leiden cluster assignments. (C) Microglia plotted by UMAP embedding showing spectral protein counts from spectral flow cytometry data for microglial phenotypic markers. (D) Violin plots annotated by microglial cell sub-cluster classification based on protein expression quantifying eight microglial subtypes across groups. Violin plots represent mean ± quartiles. Statistical significance among groups were assessed by two-way ANOVA with Fishers Least Significant Difference post-hoc test with p-values indicated above each relevant comparison. N=4/group for VaD groups and N=5/group for control groups. n=140,296 microglia.

**Figure S3. Spectral Flow Cytometry Validates Murine Brain T cell Memory Accumulation with Age**

(A) Gating strategy for identifying live singlet T cells as CD45^+^ CD8a^+^ or CD4^+^ cells. This population was then analyzed using principal components analysis (PCA), uniform manifold approximation projection (UMAP), and Leiden clustering. (B) T cells shown by UMAP embedding split by age and treatment depicting Leiden cluster assignments. (C) T cells plotted by UMAP embedding showing spectral protein counts from spectral flow cytometry data for T cell phenotypic markers. (D) Violin plots annotated by T cell sub-cluster classification based on protein expression represent mean ± quartiles. Statistical significance among groups were assessed by two-way ANOVA with Fishers Least Significant Difference post-hoc test with p-values indicated above each relevant comparison. N=4/group for VaD groups and N=5/group for control groups. n=69,615 T cells.

**Figure S4. Age- and Vascular Dementia-Associated Remodeling of Macrophage Compartment**

(A) Unsupervised Leiden sub-clustering of myeloid subsets resolved 9 transcriptionally distinct myeloid cell clusters. (B) Distribution of myeloid cell clusters across the 4 experimental groups (4-month Control, 4-month VaD, 22-month Control, 22-month VaD) plotted by UMAP embedding. Volcano plots identifying down- and up-regulated differentially expressed genes (FDR<0.05) in Macrophages_1 (C), Macrophages_2 (F), and Macrophages_3 (I). Pathway analysis showing enriched gene ontology (GO) biological processes (BP) and reactome pathways for Macrophages_1 (D-E), Macrophages_2 (G-H), and Macrophages_3 (J-K). N=3 biological replicates with 4-5 brains pooled per biological replicate for each group (total=52 brains). n=7,913 myeloid cells.

**Figure S5. Age- and Disease-Associated Remodeling of B cell Compartment**

(A) Unsupervised Leiden sub-clustering of murine brain SPLiT-seq B cell subsets resolved 4 transcriptionally distinct B cell clusters. (B) Distribution of B cell clusters across the 4 experimental groups (4-month Control, 4-month VaD, 21-month Control, 22-month VaD). (C) Dot plot illustrating marker gene expression across age and treatment groups. Dot size corresponds to the percentage of cells expressing the gene within each cluster. Color intensity reflects the mean normalized expression level (log-transformed counts). (D) Volcano plot identifying down-and up-regulated differentially expressed genes (FDR<0.05) in all B cells comparing aged and young (control and VaD combined) highlights genes enriched in aged brain (right side) and in young brain (left side). N=3 biological replicates with 4-5 brains pooled per biological replicate for each group (total=52 brains). n=631 B cells.

