## Supplementary figures and images for "High-Dimensional Single-Cell Analysis Reveals Coordinated Age-Dependent Neuroinflammatory Microglia-T cell Circuits in the Brain"

### Supplemental Figures

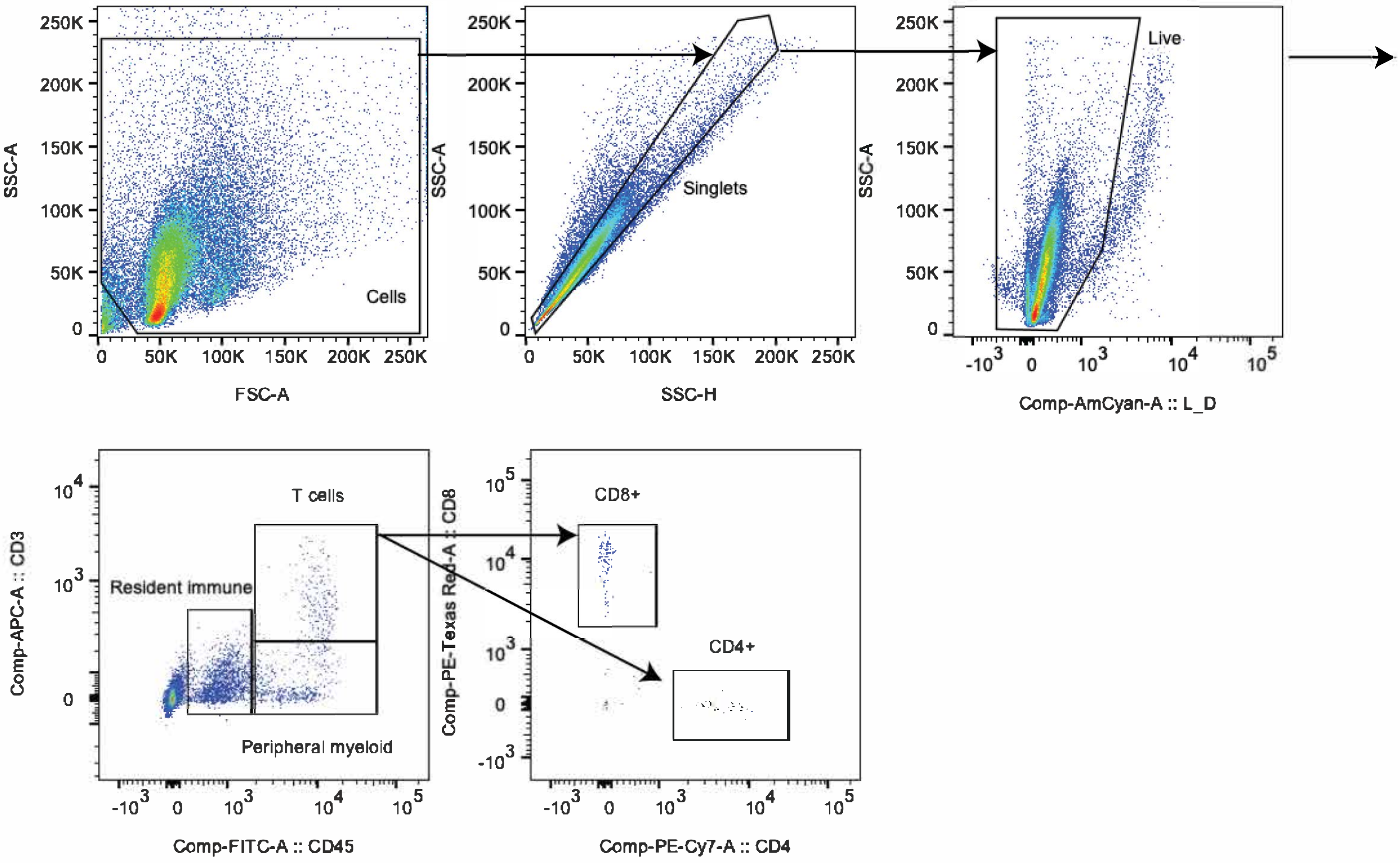

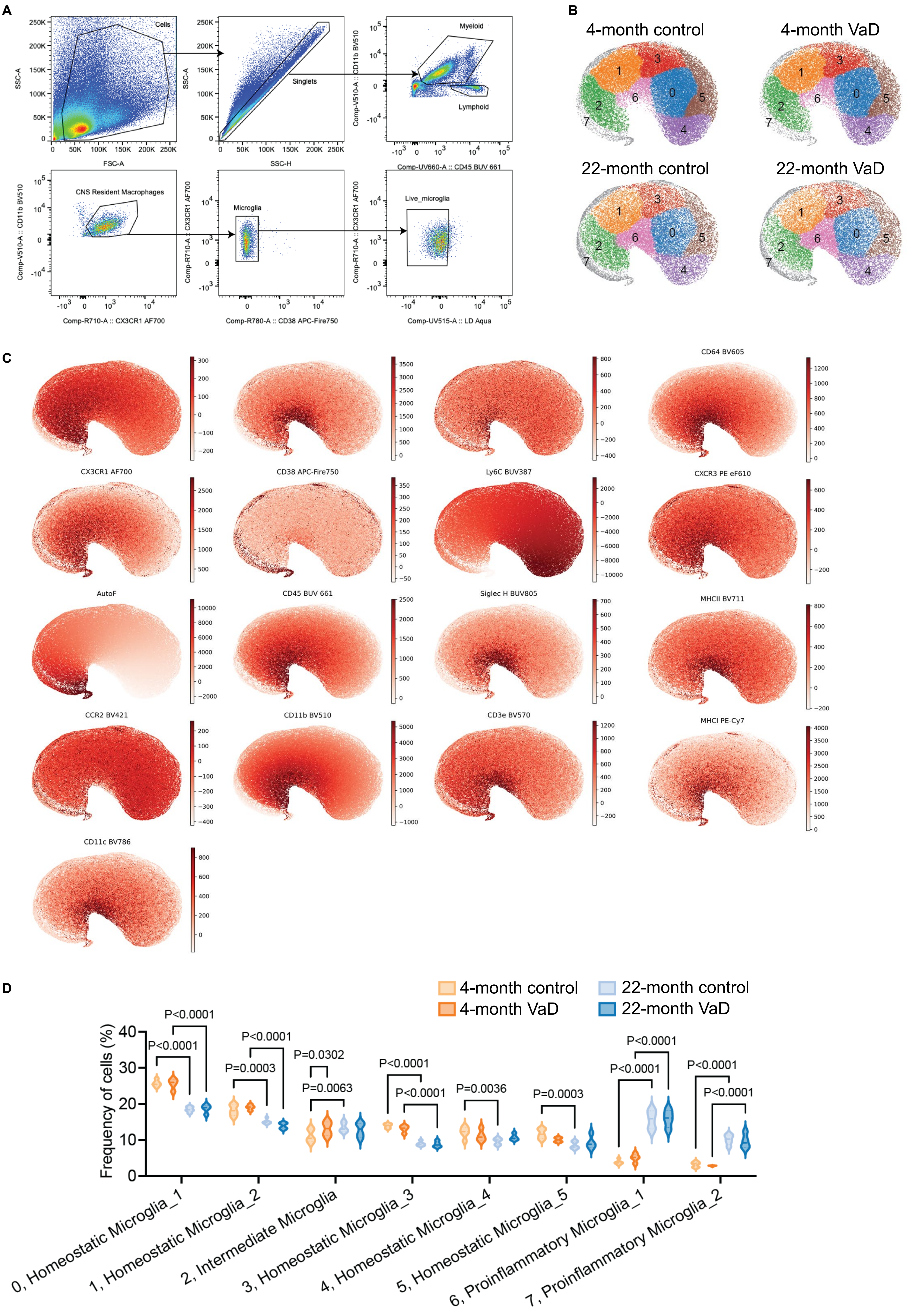

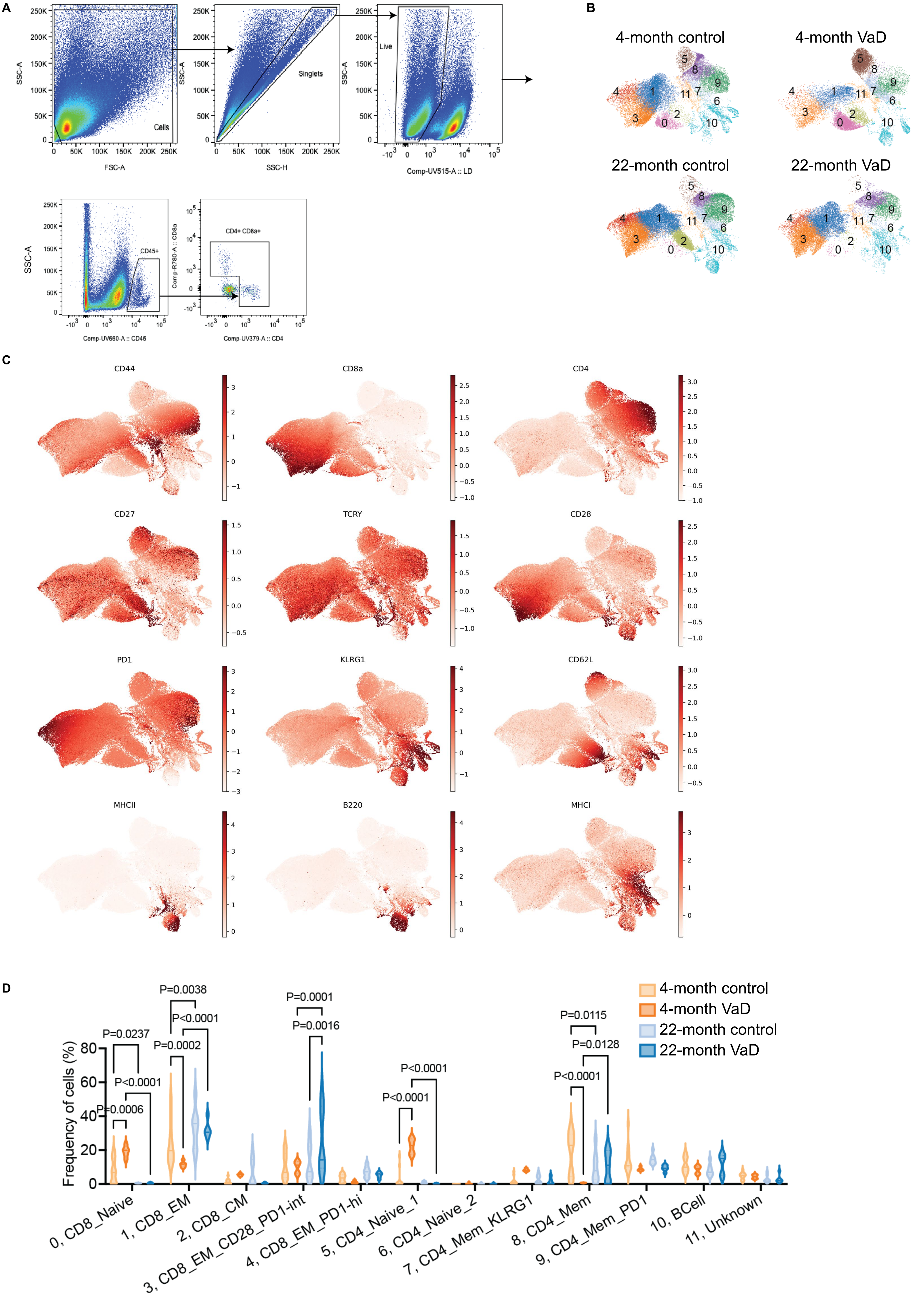

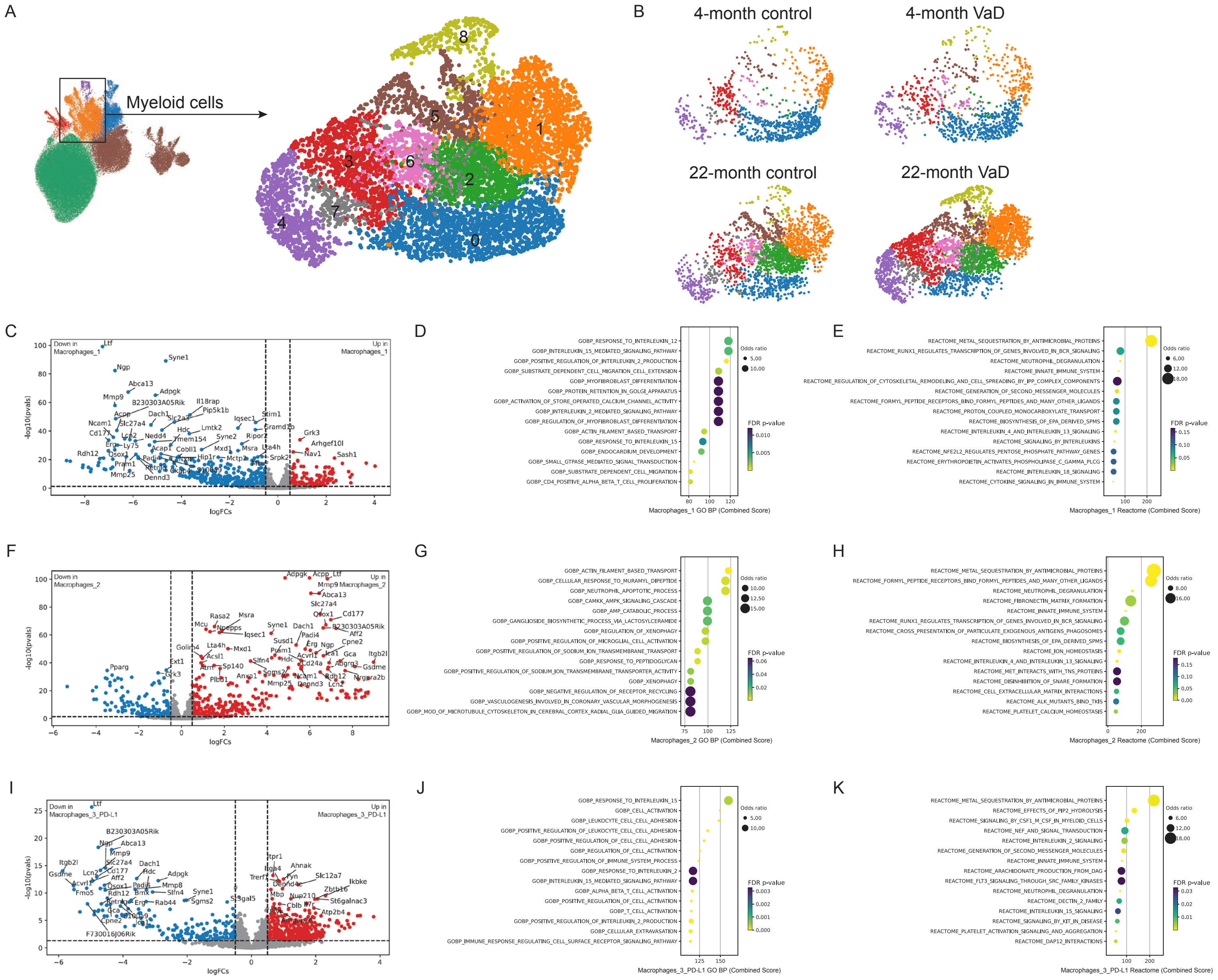

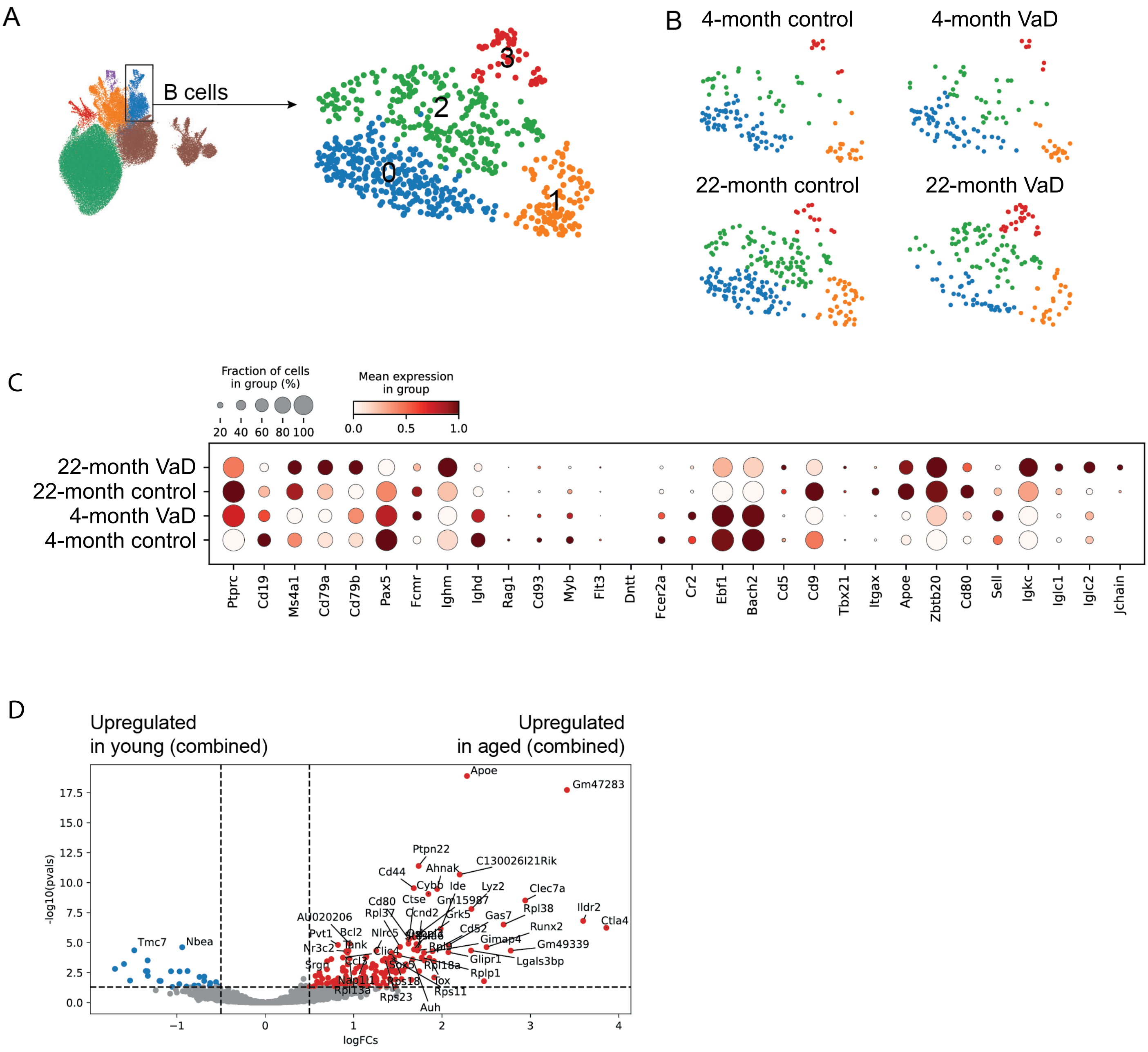
